# Decoding the Mechanism of Specific RNA Targeting by Ribosomal Methytransferases

**DOI:** 10.1101/2021.09.15.460497

**Authors:** Juhi Singh, Rahul Raina, Kutti R. Vinothkumar, Ruchi Anand

## Abstract

Methylation of specific nucleotides is integral for ribosomal biogenesis and serves as a common way to confer antibiotic resistance by pathogenic bacteria. Here, by determining the high-resolution structure of 30S-KsgA by cryo-EM, a state was captured, where KsgA juxtaposes between helices h44 and h45, separating them, thereby enabling remodeling of the surrounded rRNA and allowing the cognate site to enter the methylation pocket. With the structure as a guide, factors that direct the enzyme to its cognate site with high fidelity were unearthed by creating several mutant versions of the ribosomes, where interacting bases in the catalytic helix h45 and surrounding helices h44, h24, and h27 were mutated and evaluated for their methylation efficiency. The biochemical studies delineated specificity hotspots that enable KsgA to achieve an induced fit. This study enables the identification of distal exclusive allosteric pocket and other divergent structural elements in each rMTase, which can be exploited to develop strategies to reverse methylation, mediated drug resistance.

## Introduction

Ribosomes, the central protein synthesis machinery serves as a prime target for several antibiotics that bind to various positions within it, thereby stalling one of the many steps in translation (Arenz & Wilson, 2016, Kavčič, Tkačik et al., 2020, Wilson, 2014). Naturally, pathogens have developed several resistance mechanisms to evade the inhibitory action of these drugs, leading to the evolution of drug resistance (Alekshun & Levy, 2007, Blair, Webber et al., 2015, Davies & Davies, 2010, Wozniak & Waldor, 2010). One prevalent mechanism for antibiotic resistance is by covalent modification of the ribosomal bases at specific positions that inhibit the binding of these drugs. Acetylation and methylation of conserved bases within the ribosome by specific enzymes in pathogens are the most common mode for developing resistance (Ahmed, Sony et al., 2020, Luthra, Rominski et al., 2018, Schaenzer & Wright, 2020).

Strikingly, the pathogenic methylases such as erythromycin resistance methyltransferase (Erm) that induces resistance to erythromycin exclusively methylates at position A2058 (*Escherichia coli* numbering) in the 50S subunit and aminoglycoside resistance methyltransferase (NpmA, ArmA) that modify A1408 and G1405 respectively in the 30S subunit, exhibit evolutionary similarity with the ribosomal methylases (rMTases) that are essential for biogenesis (Davis & Williamson, 2017, Husain, Obranic et al., 2011, Husain, Tkaczuk et al., 2010, Sergiev, Aleksashin et al., 2018, Svetlov, Syroegin et al., 2021). This is in accordance with the fact that rMTases perform the important job of ensuring optimal folding of the rRNA in the ribosomes, they have also evolved to have high specificity, and perhaps that is one of the primary reasons why methylation has been chosen as a prevalent resistance modification by pathogens (Davis & Williamson, 2017, Park, Kim et al., 2010). For instance, KsgA, which is involved in ribosome biogenesis, dimethylates A1518 and A1519 in the 16S rRNA, near the decoding center within the 30S ribosomal subunit, exhibits both structural and sequence similarity to the pathogenic Erm MTase. Both these enzymes harbor a conserved S-adenosine methionine (SAM) binding Rossman fold and carry out N6 methylation of adenine but catalyze disparate substrates (Bussiere, Muchmore et al., 1998, O’Farrell, Scarsdale et al., 2004). For example, Erm rMTase is able to methylate short RNA stretches but unable to act on the assembled 50S subunit (Champney, Chittum et al., 2003, Hansen, Lobedanz et al., 2011). In contrast, KsgA does not methylate short RNAs but acts only on the 30S subunit (Desai & Rife, 2006). It has been recently shown that a chimeric version of KsgA, where certain structural elements from Erm were introduced into the KsgA scaffold, altered the reaction profile of KsgA, allowing it to act on a short Erm substrate (Bhujbalrao & Anand, 2019). Thus, the ability of rMTases to recognize their cognate substrates is encoded within the enzyme and how they achieve this is an interesting question that remains to be addressed.

In order to further understand the mechanism of substrate specificity and targeting and the molecular details of N6 adenosine methylation, KsgA was chosen as a target. Its deletion cause bacteria to become resistant to aminoglycoside kasugamycin (Schluenzen, Takemoto et al., 2006). Furthermore, KsgA deletion strains exhibit decoding errors during elongation and initiation from non-AUG codon (Schluenzen et al., 2006, Schuwirth, Day et al., 2006). In addition, the *ksgA* knockout strains manifest cold sensitive phenotype and other 16S rRNA processing defects (Connolly, Rife et al., 2008). A cryo-EM structure of 30S subunit in complex with *Escherichia coli* (*E.coli*) KsgA was reported to 13 Å and the high-resolution structure of the same complex was reported very recently (Boehringer, O’Farrell et al., 2012, Stephan, Ries et al., 2021). This structure captured the complex in a state where h45 is anchored into KsgA with one of the target bases A1519, and the terminal tetraloop G1516 being flipped-out interacting with the enzyme (Stephan et al., 2021). However, details as to how KsgA inserts itself onto the 30S platform remained elusive. Here, we have determined the high-resolution Cryo-EM structure of the *Thermus thermophilus* (Tt) immature 30S ribosomal subunit in complex with KsgA to an overall resolution of ~3.17 Å. Extensive classification of the data allowed us to separate out a unique state of the ribosome in complex with KsgA in which we were able to visualize the mode of recognition of KsgA. Especially how it inserts itself between helices h44 and h45 and positions the catalytic helix h45 into the active site pocket. This structural insight is paramount in understanding the stages involved in the entry and recognition of KsgA to the 30S ribosome. Furthermore, to understand the mechanism of substrate selection by KsgA and to discern the targeting determinants, using the structure as a guide, we designed multiple mutant versions of the ribosomal smaller subunit, containing mutations both in the catalytic helix h45 and peripheral interacting helices (h27, h24, and h44) and investigated their methylation potential. Methylation activity on these site-directed mutants revealed the importance of distant tertiary interactions in regulating enzymatic activity. The structural and biochemical data allows to determine the general strategies adopted by rRNA MTases for accurate methylation and further provide insights to develop allosteric inhibitors of rMTases that mediate antibiotic resistance.

## Results

### Cryo-EM maps of KsgA and the small ribosomal subunit

The map of the KsgA-30S complex from the full data set showed heterogeneity (Fig S2) and subsequent 3D classification yielded five main classes (K1, K2, K4, K5, and K6) with KsgA bound to the ribosomes and the overall resolutions of the cryo-EM maps range between 3.17 to 3.6 Å (Fig S2, S3, S4, and Table S1). It was observed that class K1 (sub-classified into two classes K1-k2, K1-k4) and class K6 had density for a second KsgA monomer near helix h16. Analysis suggests that this is an artifact and binding at this site is due to usage of 20-fold excess of KsgA during preparation of the 30S-KsgA complex and is not a functionally relevant site (Fig S5). Moreover, sub-class K1-k2 has a very weak map for KsgA at the 30S platform and there are no perturbations in the central decoding region (CDR) and h44 observed (Fig S6). The h28 connecting neck region between the head and body along with the loop connecting h44 and h45, which are in generally disordered, were clearly visible in the map and could be easily modelled (Fig S6 and S7). The class (K1-k2) probably mimics the stage where KsgA is scanning the binding region on the platform with its head domain before latching on the 30S, with its positively charged residues. Classes K2, K5, and K6 have h44 in a disordered state and KsgA is bound to the platform region of the 30S ribosome (Fig S2 & S7). This might depict the first state where KsgA docking onto the 30S ribosomal platform surface, increases the flexibility of h44 to accommodate KsgA. Class 4 and sub class K1-k4 with ordered h44 shows KsgA bound on the platform with its head docked over the negatively charged platform region formed by the interaction of h27, h24, h44, and h45 loop (Fig 1).

**Figure 1.**
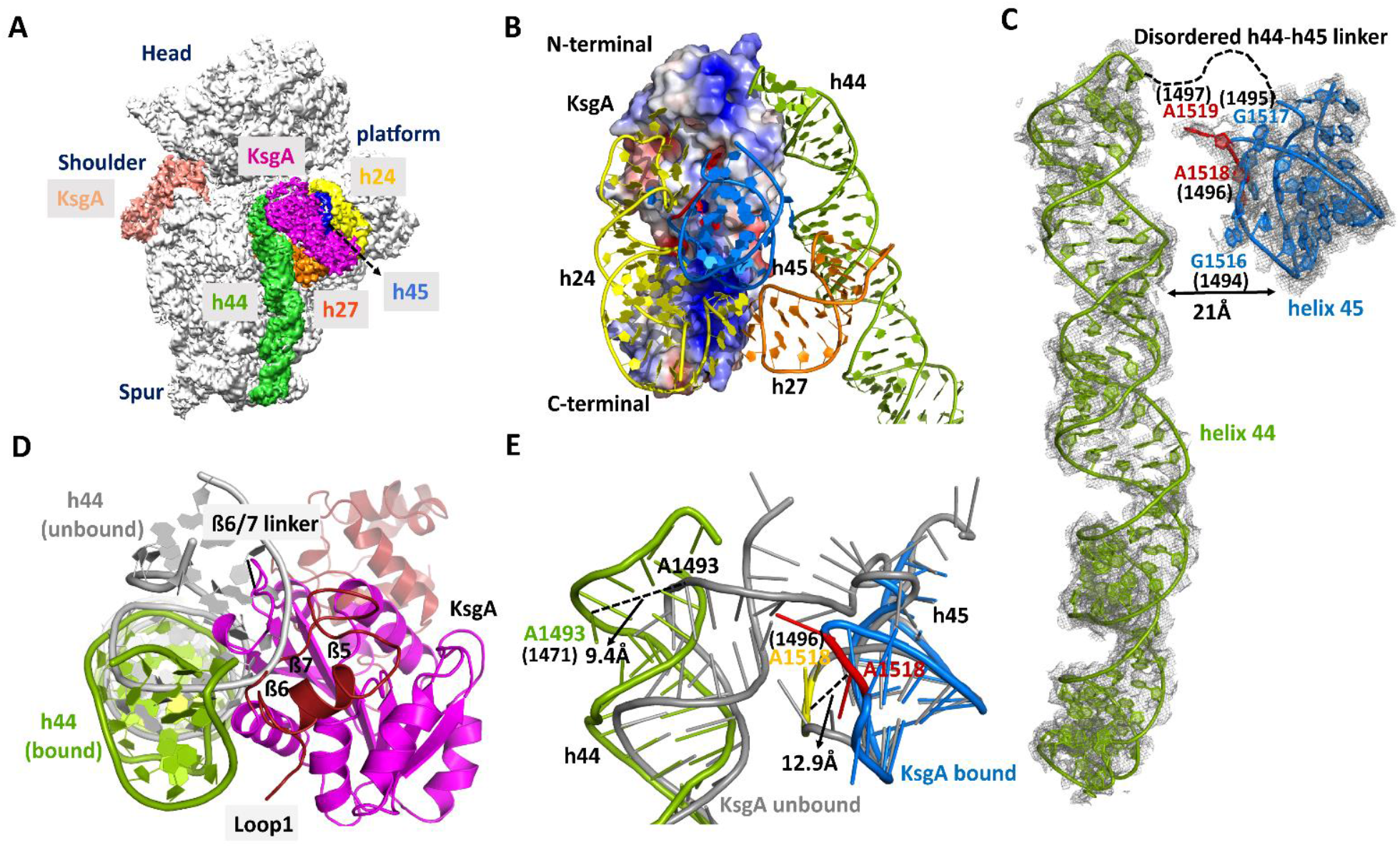
Cryo-EM structure of the KsgA-30S Complex. **(A)** Cryo-EM reconstruction corresponding to K1-k4 class, KsgA (magenta) binds to the platform and is surrounded by rRNA: h45 (blue), h27 (orange), h24 (yellow) and the upper region (green) of h44. **(B)** Electrostatic surface potential map of KsgA colored by surface charge (calculated with Adaptive Poisson – Boltzmann Solver, the scale ranges from −5kT/e (red) to 5kT/e (blue)) in complex with h45, h44, h24 and h27. **(C)** Cryo-EM densities of h44 and h45 showing the arrangement of helices in 30S-KsgA complex. **(D)** Depicts the displacement of h44 compared to unbound state (gray) verses bound (green) to accommodate KsgA β6/7 linker and loop1 onto the 30S platform. KsgA is shown in magenta with highlighted loop1 and C-domain in firebrick red. **(E)** In order to present the target residue (red) to KsgA the displacement of h44 and h45 is depicted. The unbound 30S h44 and h45 helices are shown in gray (PDB ID: 3OTO) whereas, the bound form of h45 and h45 are shown in green and blue, respectively (numbering shown in bracket is for Tt). To clearly visualize the ribosomal structural rearrangement, the model of KsgA is not shown.

Superposition with the recently published *E. coli* 30S-KsgA structure reveals that the binding mode observed in our studies is congruous to the cryo-EM reconstructions (Boehringer et al., 2012, Stephan et al., 2021) (Fig S8). For the structures reported in this study, density for the 30S head region is more pronounced in all the classes (Fig S3 & S4) and the comparison of the various populations shows that the head region exhibits heterogeneity and its swiveling movements are attributable to h28 (neck) flexibility (Fig S9).

### Role of h44 in recognition of KsgA

Helix 44 is one of the most important helices involved in decoding and harbors two bases at 1492 and 1493 (*E. coli* numbering) that scan for the correct anticodon-codon sequences (Yoshizawa, Fourmy et al., 1999). In the h44 ordered structure, substantial conformational changes were observed upon binding of KsgA with the 30S, which opens up the canonical engaged conformation of h44 and h45 to a conformation where they are separated by more than 20 Å with respect to the unmethylated immature ribosomal structure (Fig 1 C). The flexibility of h44 in the immature 30S subunit is possibly one of the reasons for the inability to trap this state in previously reported structures. Similar flexibility in the h44-45 linker is also observed in the low resolution cryo-EM structure of the eukaryotic pre-40S-assembly factor complex where the human homolog of KsgA, Dim1 was found bound as one of the proteins (Johnson, Ghalei et al., 2017, Scaiola, Peña et al., 2018).

The high-resolution structure of the 30S-KsgA complex reveals that distortions in both h44 and h45 are introduced by steric interaction of the N-terminal β6/7 linker of KsgA with h44-45 linker, in order to present the target residue to the active pocket of the enzyme (Fig 1D, E). In this conformation, h44 has bent away from the 30S subunit platform and as a result, the decoding site, importantly A1492/A1493 and C1054 moves away, thereby precluding the interaction with mRNA/tRNA duplex. Comparison of the unmethylated and methylated versions of the 30S reveals that dimethylation at h45 by KsgA results in closure of the gap between h45 and h44 that finally orders h44, maturing the 30S ribosomal subunit (Fig 2). Thus, before methylation, h44 is largely flexible and allows entry of KsgA (Fig 2, stage 2). The KsgA inserts itself between h44 and h45, separating these helices, and gets anchored onto the immature 30S through its C-terminal domain, which has several positively charged residues (Fig 1B, Fig 2, stage 3, Fig S6). This enables the interaction of KsgA and position the correct substrate, h45, into the catalytic center via flipping of the G1516 into the exclusive pocket prepping the catalytic pocket for methylation by allowing flip of A1518/A1519 (Fig 3 A, B, C). Subsequently, methylation mediated volume changes at the target adenosine bases triggers the release of KsgA from the 30S platform. In the final methylated form, both helices h44 and h45 come closer and are stabilized by both hydrogen bonding and Vander Waal interactions thereby, sealing the interface that results in a structured h44 such that the biogenesis is complete and the ribosome is primed for further assembly (Fig 2, stage 4).

**Figure 2.**
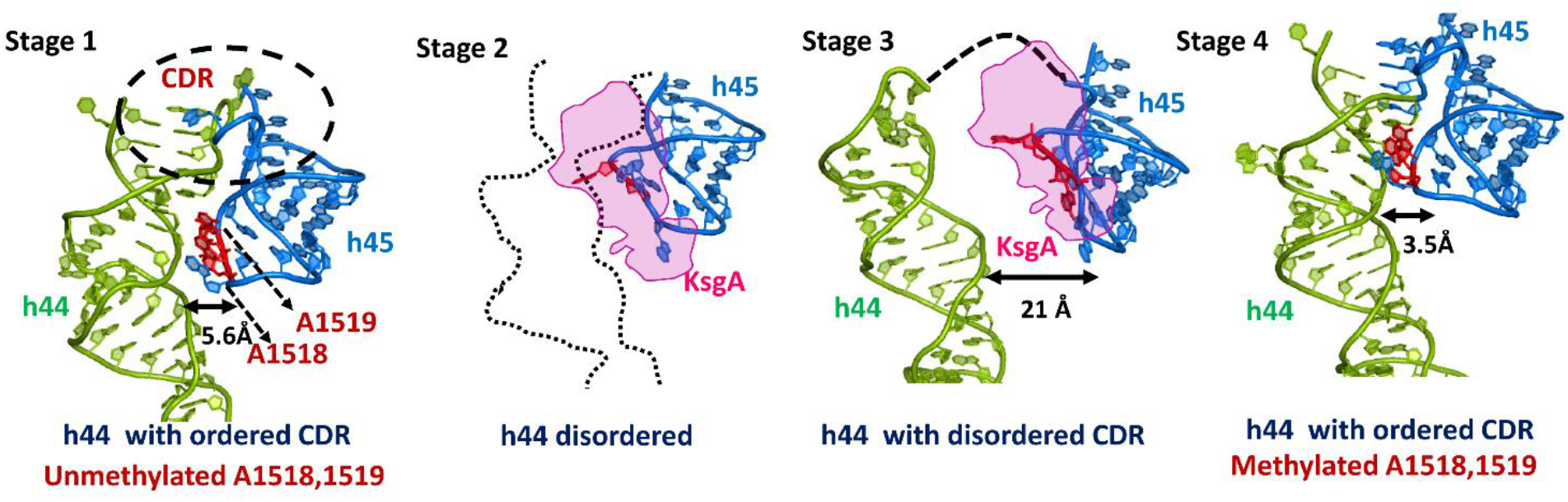
Steps involved in KsgA methylation during biogenesis to achieve mature 30S. Stage1 is the pre-methylated stage where h44 (green) and h45 are loosely packed. At Stage 2, h44 adopts a flexible conformation (shown as dotted line) allowing entry of KsgA (magenta) (Similar to the state observed by Stephan *et al*.) This later rearranges in Stage 3 to a conformation where h45 (marine blue), presents A1518 and A1519 (red) to the KsgA catalytic pocket for methylation. In Stage 4, methylation leads to dissociation of KsgA and orders the h44 and h45 conformation so as to achieve an active central decoding center (CDR). (PDB ID: 3OTO (unmethylated A1518/19, Stage1), 4B3R (methylated A1518/19, Stage4). Stage 2 (K5) and Stage 3 (K1-k4) are experimentally observed as cryo-EM maps in this work.

**Figure 3.**
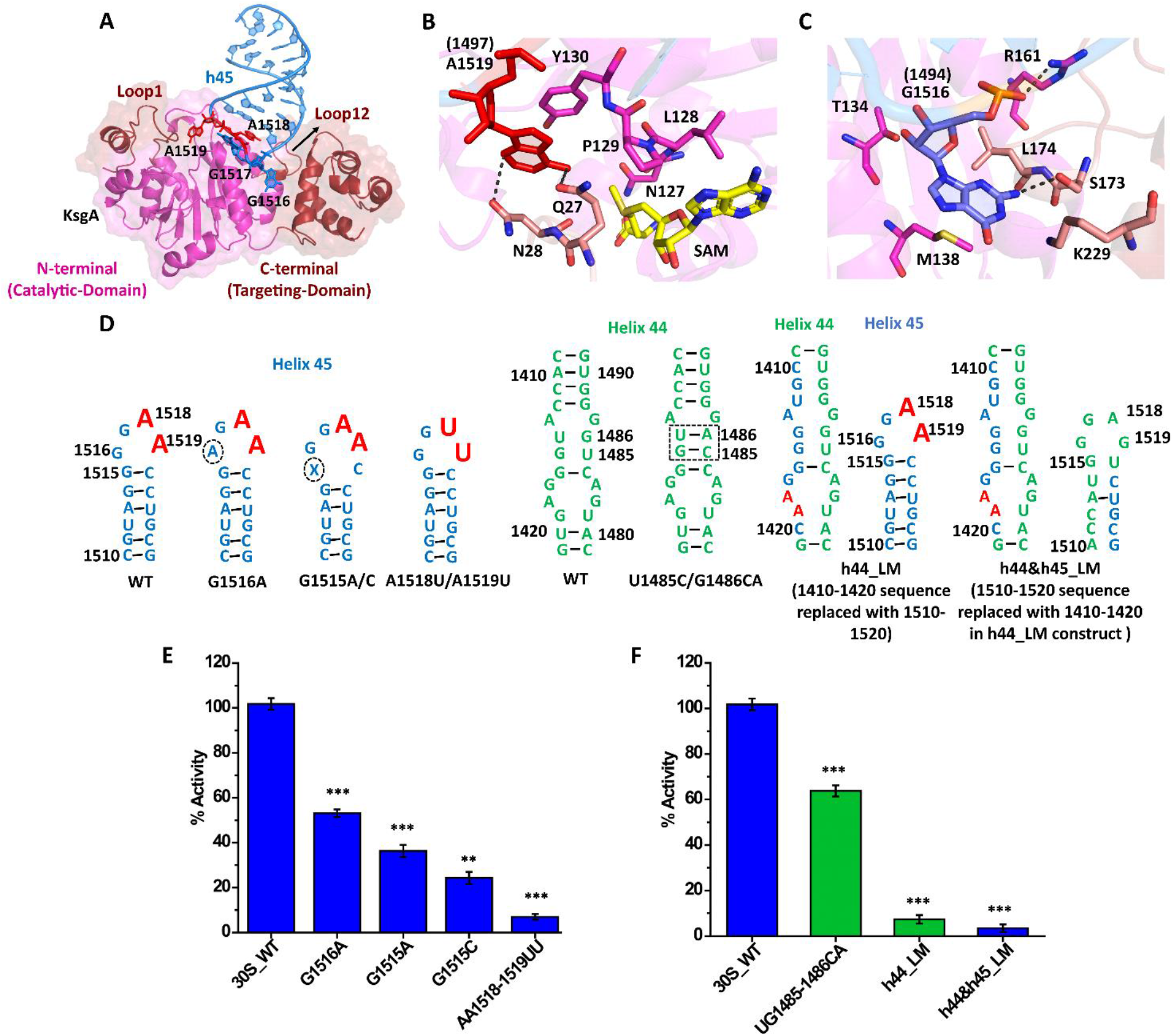
Effect of mutagenesis of select bases in h45 and h44 of 30S present near the KsgA interaction region. **(A)** Schematic diagram depicting the interaction of GGAA tetraloop of h45 and KsgA. **(B)** The active site of KsgA with key residues shown in stick representation, pocket with flipped A1519 (red) (numbering shown in bracket is for Tt) into catalytic pocket, carbon atoms of the interacting residue are shown in magenta and salmon and those of SAM in yellow while oxygen and nitrogen atoms are colored in red and blue respectively. SAM is modelled by using PDB ID: 6IFT, not present in 30S-KsgA complex structure. **(C)** Interaction of G1516 (blue) (numbering shown in bracket is for Tt) with KsgA via a pocket formed at the interface of its N and C-terminal domain. Loop12, C-terminal domain residues are shown in salmon and N-terminal residues are shown in magenta. **(D)** Design of h45 and h44 mutants, KsgA target residue are highlighted in red, residues encircled are subjected to mutagenesis. **(E)** In-vitro KsgA methylation assay with constructs harboring mutation in h45 tetraloop. **(F)** In-vitro KsgA methylation assay with altered target site constructs of h44. Data for three experimental replicates are shown with the mean and standard deviation. Histogram colors are as per the scheme chosen for each 30S helix in Figure1. Student’s t-tests was used to calculate P-values (**P<0.01, ***P<0.001) for the comparisons of 30S_WT (wild type) with 30S mutant variant. Results show that methylation efficiency is significantly affected on perturbation of these important regions.

### Mechanism of substrate fidelity of KsgA

The cryo-EM maps and models reveal that for productive methylation of the two adenines, KsgA positions itself by establishing extensive interactions with the 16S rRNA formed via apposition of helices h27, h24, h45, and h44 (Fig 1 B). Although these four helices are distantly placed in secondary structure but upon folding and assembly in the ribosomes, these helices come close together to form a complex interface recognized by KsgA (Fig 1 A, B). These rRNA helices surround KsgA and this extensive binding mode clearly supports the rationale for requirement of fully assembled immature 30S rather than a short fragment of RNA or 16SrRNA as a substrate. The conserved h45 ‘GGAA’ tetraloop predominantly mediates the interaction of the ribosome with the N-terminal domain of KsgA by bending and inducing a 12 Å shift from its unbound form (Fig 1 E), such that it positions A1519 into the active site cavity. The complexation results in the flipping out of A1519 (Fig 3 A, B, S6), a conformation commonly observed in several MTases and necessary to facilitate catalysis (Jiang, Li et al., 2017, Lee, Agarwalla et al., 2005, Lin, Chen et al., 2020, Liu, Long et al., 2017, Ma, Ma et al., 2016).

Comparison of the 30S apo structure with the 30S-KsgA complex further shows that entry of KsgA pushes h44 inwards and its upper portion swings by ~ 9.4 Å to accommodate the linker that connects β6-β7 as well as loop1 and loop12 of KsgA (Fig 1 D, E). This rearrangement is in accordance with previous reports from chemical probing studies, where this region of h44 is protected in the presence of KsgA (Xu, O’Farrell et al., 2008). Thus, in order to present the conserved adenine to the rMTase active site, displacement of both h44 and h45 from the canonical conformation occurs. Furthermore, to anchor itself to the ribosome, KsgA undergoes a substantial change in the SAM binding loop1 (rmsd 5.0 Å, between 30 N-terminal amino acids), the interface loop12 (rmsd 1.27 Å, between 13 amino acids) as well as C-terminal domain (rmsd 2.48Å, between 75 C-terminal amino-acids). Several positively charged residues contribute to the interaction interface via contact with the stem loop of h45 (1513-1515) (Fig S10 A). In addition, G1516 inserts into a shallow cleft that lies at the interface of the head and loop12 in KsgA (Fig 3 A, C, S6). Comparison with the published structure by Stephan *et al*. reasserts that G1516 region is indeed important for positioning of the target RNA (Stephan et al., 2021).

To further understand the importance of base conservation in stabilization of the tetra-loop and its sensitivity toward perturbation to catalysis, we constructed several variant versions of the ribosome. As a negative control, we mutated A1518, A1519 to uracil and as expected, methylation activity was obliterated, and no incorporation of tritium labeled SAM was observed (Fig 3E). Mutating the other important interacting tetraloop base G1516 to an alternate purine base, adenosine, resulted in 50% loss in activity (Fig 3 D & E), iterating the importance of the correct base stabilization in the loop12-C-terminal cleft of KsgA. We also find that when the C-terminal domain of KsgA is deleted (ΔC-KsgA), the activity is reduced close to 50% of the native enzyme, which is similar to the G1516A mutant (Fig 4D). This is because in ΔC-KsgA the cleft is not fully formed resulting in incomplete stabilization of G1516.

**Figure 4.**
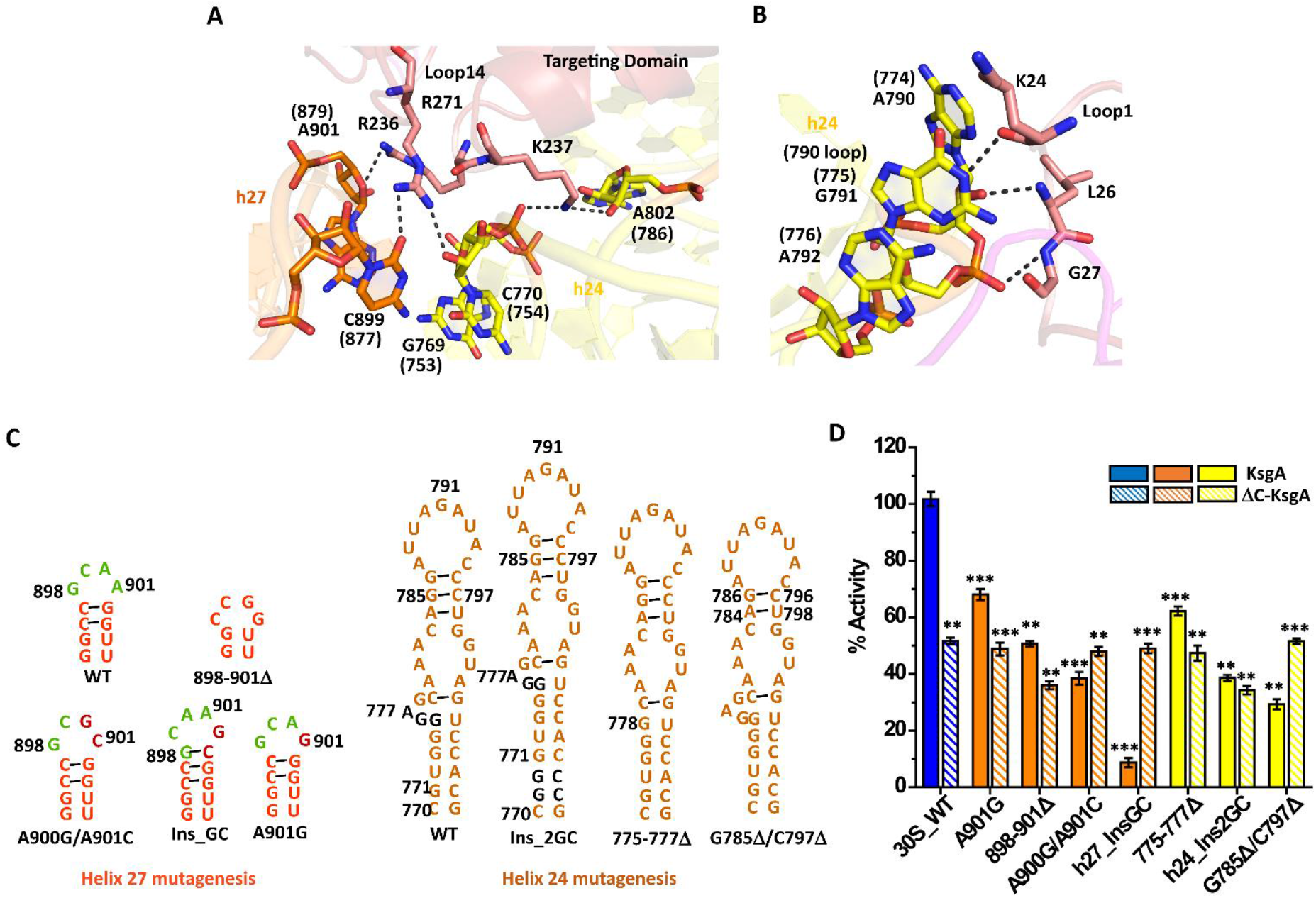
Probing tertiary interactions of KsgA with h24 and h27. **(A)** The highly basic loop14 of KsgA targeting domain recognizes the juxtaposition of helix 27 and 24 through multiple interaction with rRNA backbone (numbering shown in bracket is for Tt). **(B)** Loop1 interaction with helix 24 (790-loop) phosphate backbone (numbering shown in bracket is for Tt). The density for the residues 773-779 is poor in all classes with KsgA bound and not modelled. In this figure, using the K1-k2 model, this region has been extrapolated to show potential interaction built on K1-k4 map. **(C)** Design of h27 (GCAA, green, a conserved 900-loop) and h24 mutants. The residues that were mutated/inserted/deleted are highlighted in maroon and black for h27 and h24 respectively. **(D)** In-vitro methylation assay using 3H-SAM with native and ΔC-KsgA. Histogram colors are as per the scheme chosen for each 30S helix in Figure1. Data for three experimental replicates are shown with the mean and standard deviation. Student’s t-tests was used to calculate P-values (**P<0.01, ***P<0.001) for the comparisons of 30S_WT (wild type) with 30S mutant variant.

The terminal tetraloop bases in RNA tetraloop recognizing enzymes have been shown to be important in establishing binding surface with the protein and any mutation in these bases results in significant decrease in the binding affinity (Thapar, Denmon et al., 2014). The base G1516 is ~15 Å away from the main catalytic pocket and yet has a marked effect on activity, signifying that long distance allostery is at play. When an adjacent base in the tetraloop, G1515 is mutated, either to adenine or cytosine, these ribosomal mutants exhibit 60% and 80% reduced activity respectively, (Fig 3D & E), strengthening the importance that the stem region adjacent to the catalytic tetraloop not only plays an important role in interacting with KsgA but also for stabilization of the tetraloop conformation itself. The significant loss of activity observed for the G1515C mutant possibly results due to the disruption of the Watson-Crick base pairing interactions at this site and results in widening of the tetra loop that subsequently results in disruption of the overall interactions of h45 with KsgA (Fig 3D & E). Note that in the Tt rRNA, the equivalent residue of G1515 is C1493 and it base pairs with G1498.

### Mutation of the h44 region and shuffling of the target site

Helix h44 of 16S rRNA contacts KsgA via a distorted bulge (1485-1486), an interaction responsible for h44 to pivot towards KsgA. Thus, we asked if the ability of methylation is affected with the removal of the bulge. The bulge was replaced with a perfect Watson-Crick match at this position and it was observed that the activity was reduced by 40% (Fig 3D & F) showing that loss of anchoring at this position does affect methylation efficiency. Previous chemical probing experiments have shown that this area is protected in the presence of KsgA (Xu et al., 2008), which is not surprising as the structure shows that the region 1482-1497 of h44 indeed comes close to KsgA (Fig S10 C) forming an interaction interface. We conclude that this region is important for both, interaction with KsgA and to allow flexibility to the h44 during methylation by KsgA.

The above observations raise the question if the tetraloop architecture is sufficient or the surrounding h44-h45 conformation is also essential for methylation. To address this, the tetraloop and stem region were stappled at other positions within the ribosome and methylation efficiency was gauged. Chimeric ribosomes where a portion of h44 was replaced by h45 such that the ribosome had two catalytic tetraloops was created and it was found to be completely inactive (Fig 3D & F). In the absence of properly positioned h44 and h45, the ribosome is unable to support proper binding of KsgA at either of the two catalytic centers. 3D RNA composer predicts that the replacement of h44 region by h45 completely melts the overall structure of h44, which then changes the orientation of h45 away from the canonical conformation resulting in a different 3D architecture (Fig S10 D) that is not recognized by KsgA. The fact that this shuffling (h44-LM) has resulted in the reorganization of the h44-h45 interface is supported by the non-viability of Δ7prrn strain as oppose to other minor variant ribosomes (Table S2). Another chimera was created where h45 and h44 interacting region was swapped. However, this shuffled ribosome also showed no activity suggesting the order and appropriate placement of peripheral regions are paramount for the efficient functioning of KsgA. In both cases, production of ribosome was confirmed by sucrose gradient in the MS2 tagged system, which harbor both plasmid born tagged and intrinsic untagged ribosomes (see Material and Methods). In summary, the shuffling studies strengthen the idea that KsgA recognizes the mini-tetraloop structure in a complex environment with correctly placed h44 and h45 and other surrounding helices to enable the catalytic conformation, explaining why it does not methylate mini-RNA templates.

### Interaction of KsgA with peripheral helices h27, h24

As mentioned earlier KsgA interacts with other important helices such as the h27 switch and h24 that are present near the decoding site. These helices along with h44 rearrange during biogenesis to create a conducive environment for binding of the initiation factors as well as 50S such that the catalytic machinery can be assembled for protein synthesis (Carter, Clemons et al., 2000, Laursen, Sørensen et al., 2005). Three loop structures of KsgA, loop1 (N-terminal loop), loop12 (peripheral loop at the opposite end of N-terminal domain) and loop between α10 - α12 (loop 14) that lies in the C-terminal domain primarily interact with the peripheral rRNA. While loop12 holds both h45 and h27 on the front interface (Fig S10 B), loop14 anchors h24, and h27 tethering them via a string of positively charged interactions (Fig 4A). Here, bases C899 - A901 in h27 as well as bases G769, G770, and A802 in h24 juxtapose via a bridging interaction of loop14 residues R236-K237 (Fig 4A). Moreover, R271 from loop15 also hydrogen bonds with both the helices further strengthening the interface (Fig 4A). The highly extended conserved basic surface of loop14 seems to impart specificity of recognition to KsgA for this site. While one end of helix h24 is tethered by loop14, it extends further and runs across the entire backside of KsgA making multiple contacts (Fig 1B). More importantly, h24 makes important hydrogen bonding contacts with loop1. Loop1 acts as a lid and its opening and closing serves as an entry path for SAM. Here, bases A790 and G791 on h24 can form a potential hydrogen bonds with residues K24 as well as backbone of L26 and G27 on KsgA (Fig 4B). It has been reported earlier that mutation of K24 results in substantial loss of activity (Bhujbalrao & Anand, 2019) and therefore h24 plays an indirect role in controlling reactivity via interaction with loop1.

To access the importance of some of these peripheral interactions multiple MS2 tagged variants were constructed where specific bases were either replaced or deleted. The aim was to disrupt the interactions that hold helices h27 and h24 to KsgA. A point mutation involving A901G in h27 was performed as it contacts both loop12 and loop14. We observed 70% activity (Fig 4C, D), hinting that a purine-to-purine exchange does not alter the conformation much as most of the contacts are via the phosphate backbone interactions. On the other hand, A900G/A901C mutation has only 40% activity as changes in these bases disturbs the overall conformation of this region (Fig 4C, D). Shortening of helix h27 by creating a deletion at position 898-901 also results in 50% loss of activity (Fig 4C, D). We believe that the replacement of the bases in this region results in non-conducive interactions with the head, reiterating the importance of a C-terminal anchor point. In fact, a mutation that has an increased length via insertion of a GC pair at the position between 901-902 results in complete loss of activity for the full length KsgA (Fig 4C, D). Lengthening of this region results in a direct steric clash with the C-terminal domain of KsgA and does not allow KsgA to position itself at its cognate position. The activity of the longer h27 is restored to 50% of the native when ΔC-KsgA was used for methylation reasserting that steric clash with the head domain was the reason for loss of activity.

Mutations on h24 that run along the entire backside of KsgA were also engineered. Several mutations in order to shorten or lengthen h24 to disrupt individual contacts at the N-terminal region were made. Here, the most significant effect was on lengthening the helix h24 by two base pairs, where the methylation efficiency was reduced by 60% both with native and ΔC-KsgA (Fig 4C, D).

This is because the longer h24 again causes a steric clash with loop1 displacing it from its original position and thus likely to affect the entry and exit of SAM. A similar loss in activity of about 60% was reported when G791 which contacts loop1 via K24 was disrupted (Desai, Culver et al., 2011). To investigate the importance of the flexibility of h24, the hinge region at position GGA775-777, observed below the 790 loop of h24 which probably imparts motion was deleted and a shortened h24, G785Δ/C797Δ construct was created. It was observed that this shortened and rigid version of h24 again shows 30% and 60% of activity (Fig 4C, D) respectively. Since h24 anchors at both ends of KsgA, shortening of h24 induces strain in the helix and this shorter version is neither able to reach loop1 and help in its placement effectively nor is it able to properly engulf KsgA in this strained form. Corroborating this data, it was observed that in the ΔC-KsgA version where the C-terminal anchor point is deleted, methylation activity no longer depends on perturbation of h24 length and is always reduced to 50% of native. This data confirms that h24 and h27 help in proper positioning of the head of KsgA and enhance its methylation potential by serving as a peripheral anchor that prevents dislodging of KsgA from the ribosome.

## Discussion

### Insights into the mechanism of methylation

Ribosomal MTases whether from pathogenic organisms or those involved in biogenesis, display high fidelity and only methylate the cognate bases(Pletnev, Guseva et al., 2020, Sergiev et al., 2018). Thus, it appears that they have been programmed to accurately recognize the correct sequence and this information is embedded in their structure. Comparison of the Rossmann fold harboring MTases reveals that the substrate enters perpendicular to the body of the enzyme and inserts itself into the catalytic pocket by inducing a base flip (Fig S11). The flipping of the methylating base is a common feature observed in several other MTases including DNA MTases and seems to be a prerequisite for proper binding of the substrate (Goedecke, Pignot et al., 2001, Horton, Liebert et al., 2006, Wilson, Kellie et al., 2014). In all these cases the flipped base is stabilized in the catalytic site via π-π interaction of the adenine moiety with tyrosine or a phenylalanine residue (Fig S12 D).

One of the questions that remains a matter of debate is how do these enzymes enable dimethylation? In each round, a fresh SAM has to enter to execute the reaction therefore the entry of SAM and exit of SAH requires a substantial motion of the protein or the RNA architecture. Furthermore, in the case of KsgA successive bases are methylated thus a shift in the base that is to be methylated has to also occur. Stephan *et al.* using their recent structure of *E. coli* KsgA-30S complex as a guide proposed that G1516 is the anchor point and to achieve tandem methylation of A1518 and A1519, G1517 flips to facilitate A1518 methylation after A1519 is already dimethylated (Stephan et al., 2021). Our structure also shows that G1516 fits snuggly in the positively charged pocket (Fig 3B, S6, S8) and it appears to be one of the selectivity checks to allow for the correct RNA.

Analysis of the conformation of tetraloop in the unmethylated, versus protein bound and the post-methylated versions points to the hypothesis that it is the malleability of the tetraloop structure and the overall backbone conformations that determine the manner in which bases are presented to the active site. In presence of the protein, the mix of hydrophobic and electrostatic interactions alters the conformation of the tetraloop by a rotation of approximately 90° (Fig 5A), inducing significant distortions in the backbone such that it fits into the active site pocket and in this new conformation both G1516 and A1519 flip out and occupy tailored protein pockets. A similar scenario is observed in the ribosomal sarcin tetraloop which significantly changes conformation in presence of the protein (Thapar et al., 2014). The conformation of G1517 is most likely determined by the overall backbone conformation of the tetraloop and can vacillate between the various partially stacked states as is seen in KsgA and TFB1M complexes. An NMR structure determined by Liu *et al*. of the methylated h45 tetraloop shows that after methylation A1519 is flipped in, whereas A1518 is flipped out suggesting that tandem methylation can be facilitated via local flipping motions concentrated around the bases which undergo methylation(Liu, Shen et al., 2019). However, once h45 gets disengaged from the protein, the tetraloop backbone again undergoes a major reorganization taking a conformation, as is observed in the mature 30S, where G1516 is flipped back resulting in a concomitant flipping out of A1519 (Fig 5B). Thus, we propose for consecutive methylation after A1519 gets di-methylated it flips back towards h45 allowing A1518 to get methylated. The methylation of A1518 triggers the release of the RNA from the protein, purporting a reorganization of the tetraloop resulting in a conformation where both the methylated bases are flipped out such that they engage closely with h44 (Fig 5 B). To allow for the successive methylations it is highly unlikely that KsgA moves out of the 30S and re-enters every time, rather we propose the protein is held into position by the positively charged C-terminus. The structure shows that the C-terminal of KsgA serves as a tight anchor and holds the enzyme in position via h27 and h24 and therefore we speculate that this allows the enzyme for successive methylations without leaving the vicinity of the rRNA. This could be a common mechanism for other rMTases, where the C-terminal serves as a crucial anchor. The importance of C-terminal in imparting methylation efficiency is asserted with our data for KsgA, which shows only 50% activity in the absence of its head domain. Moreover, it has been reported that the headless Erm37 from *Mycobacterium tuberculosis* shows a mixed methylation profile and in addition, it is an error prone MTase likely due to insufficient anchoring (Madsen, Jakobsen et al., 2005).

**Figure 5.**
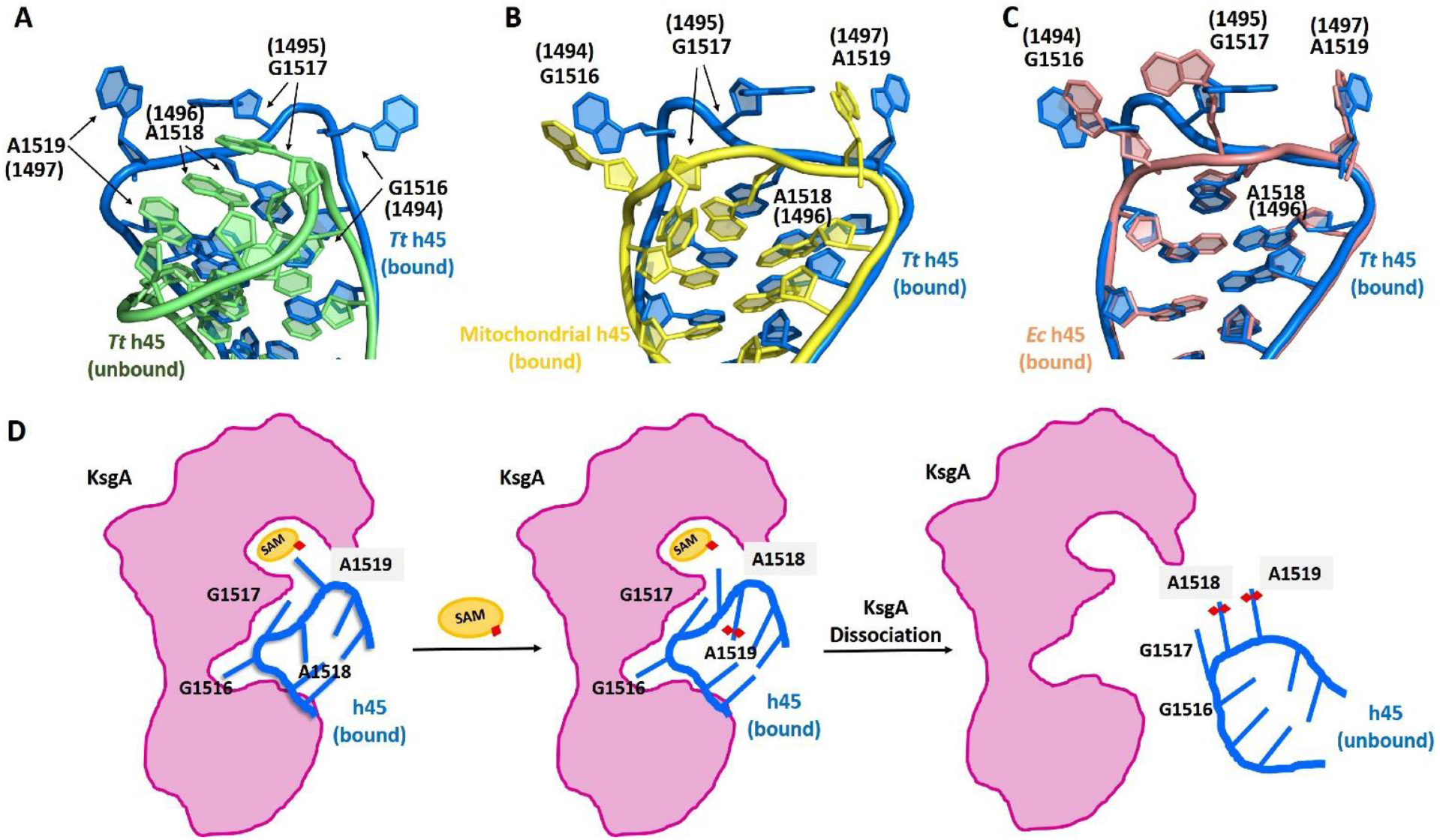
Tetraloop conformational flexibility. **(A)** An overlay of the h45 tetraloop bound to KsgA (marine blue) and unbound state (green) (PDB ID: 3OTO) of the Tt ribosome showing the conformational differences. **(B)** The conformational difference of h45 tetraloop in bound state with KsgA (marine blue) and TFB1M (yellow) (PDB ID: 6AAX). **(C)** An overlay of Tt h45 tetraloop (marine blue) in bound state with BsKsgA and Ec h45 tetraloop (salmon) with EcKsgA (PDB ID: 7O5H) (numbering shown in bracket is for Tt). **(D)** Proposed mechanism of dimethylation. Tetraloop conformational change induces A1519 flips into the active pocket. Dimethylation of A1519 results in further local flipping of adjacent A1518 base. Finally, dimethylation of A1518 dissociates the enzyme from 30S and h45 resumes its native conformation as observed in mature 30S ribosomal subunit.

### Selectivity of cognate substrate via divergent loop elements

One of the striking features is the specificity of each rMTase for a particular RNA sequence. It appears that the enzymes have developed evolutionarily diverse substrate recognition loops such as loop1, loop12, and loop14 that scan for the correct substrate (Fig 6, S12 A). These loops are different in length and amino acid composition for each MTase and act as a handle that holds the extreme end of the cognate RNA substrate. In the case of the ribosomes, these loops also interact with the peripheral RNA segments and provides an additional layer of stringency to the recognition process. For instance, in KsgA both loop12 and loop14 hold h45. As shown in Supplementary Fig. 12 B, C due to its short loops 12 and 14, binding of Erm in the KsgA site is not favorable, and it will not be able to effectively interact and stabilize h45 into the catalytic center. A closer analysis shows that these loops along with a portion of the head domain form an exclusive allosteric pocket that locks the cognate site into position. This shallow pocket in rMTases is almost always 15 Å away from the reaction center and harbors a conserved base that flips into it. A similar flip is also observed in the TFB1M RNA protein complex where G934 is approximately 15 Å from the methylating base A937 also flips(Liu et al., 2019) (Fig S11 B). This peripheral base flipping seems to be a common targeting mechanism in other RNA MTases, for instance, in the mRNA recognizing protein METTL16 multiple base flips are observed to seal the interaction interface (Fig S11 B). It is also observed that in METTL16 where mRNA is the primary substrate, loop12 extends into the RNA and holds it at multiple positions highlighting the evolutionary diversity of this loop segment (Doxtader, Wang et al., 2018). (Fig S11 A).

**Figure 6.**
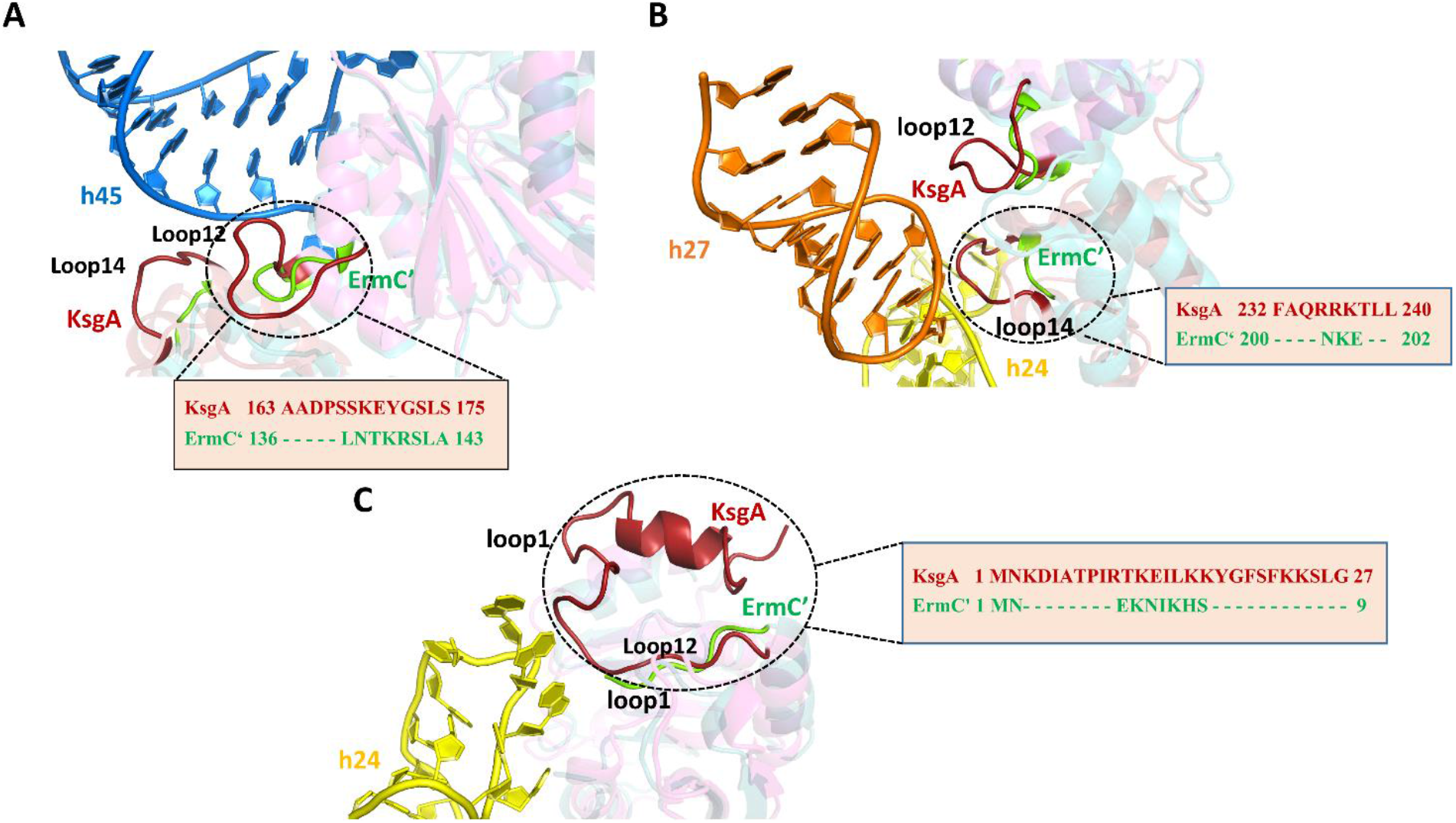
Targeting determinant of methyltransferase. **(A)** The comparison of longer loop12 and loop14 of KsgA (firebrick red) with shorter respective loops of ErmC’ (green, PDB ID: 1QAM), interacting with h45 stem region. **(B)** The comparison of longer loop12 and loop14 of KsgA (firebrick red) with shorter respective loops of ErmC’ (green, PDB ID: 1QAM), interacting with 30S platform helices h27 and h24. **(C)** Similarly, longer loop1 of KsgA (firebrick red) than ErmC’(green) interacts with h24. Respective sequence difference of KsgA and ErmC’ loops are highlighted in the inset.

The other region that plays an important role is loop1. In KsgA this loop interacts with h24 and the interaction of h24 via K24, and G791 has been shown to be a very important anchor point to maintain catalytic activity. This loop1 is found to be variable across MTases, for instance, loop1 is much shorter in ErmC’ than in KsgA (Fig 6C). Thus, it appears that depending on the position and type of RNA the MTase is accessing, the loop sizes are tailored and harbor specificity determinants. For MTase such as KsgA that targets the surface of the ribosome these loops are longer such that a correct match between the methylation site and enzyme can be achieved. However, for Erm that only methylates immature ribosome (Champney et al., 2003) the task of recognition is more to access the reaction center and hence these loops are shorter in length. To corroborate the fact that these loops indeed tune specificity, an earlier study showed that chimera’s which consisted of loop1 and loop12 of Erm onto an otherwise KsgA scaffold made it capable of methylating Erm mini-RNA substrates (Bhujbalrao & Anand, 2019).Furthermore, saturation mutagenesis of loop12 also confirmed that positively charged residues in loop12 of Erm are important for substrate recognition (Rowe, Mecaskey et al., 2020).

In conclusion, the cryo-EM structure of KsgA along with the mutagenesis studies performed here shed light on both the mechanism of targeting and recognition adopted by N6 Rossmann fold rMTases, to achieve specificity of function and additionally help in understanding the final step of biogenesis. The study reasserts that specific loop regions that are evolutionarily divergent and differ in both size and amino acid composition across MTases are the key elements that impart selective recognition of the target site. Moreover, an exclusive allosteric pocket 15 Å away from the catalytic center, at the edge of N-terminal domain and the C-terminal region acts as a selectivity filter and only allows the cognate base to properly position itself. The C-terminal region acts as a robust anchor allowing the enzyme to be properly latched onto the ribosome. These findings open avenues towards targeting allosteric sites in pathogenic MTases that can serve as selective drug design hotspots for reversing antibiotic resistance.

## Material and methods

### Purification of methyltransferase

KsgA from *Bacillus subtilis 168* was cloned in a modified pET vector as described previously (Bhujbalrao & Anand, 2019). A variant of KsgA where the C-terminal 75 amino acids are deleted (ΔC-KsgA) was also cloned into the pET vector. Both (BsKsgA and ΔC-KsgA) variant of the methyltransferase was transformed into *E.coli* BL21 (DE3) pLysS, expressed and purified according to published procedures (Bhujbalrao & Anand, 2019). Briefly, cells were grown at 37 °C until OD_600_ reached 0.6 – 0.8, and then KsgA expression was induced by adding isopropyl-β-D-thiogalactopyranoside (IPTG) to a final concentration of 1 mM and grown further for 8 hours at 25°C. The bacterial cells were harvested by centrifugation at 5000 rpm for 15 minutes and then re-suspended in lysis buffer (50 mM HEPES, pH 7.5, 3 mM imidazole, 500 mM NaCl). Cells were homogenized by a probe sonicator (Vibra-cell; SONICS, CT, USA) and the lysate was centrifuged to remove cell debris and other insoluble material. The supernatant was incubated with Ni-NTA resin pre-equilibrated with lysis buffer on a rocker for 1 hour at 4°C. Subsequently, the resin was separated by centrifugation and transferred to a gravity column. The resin was extensively washed with wash buffer (50 mM HEPES, pH 7.5, 10 mM imidazole, 500 mM NaCl) to remove non-specifically bound proteins and KsgA was eluted with elution buffer (50 mM HEPES, pH 7.5, 100 - 200 mM imidazole, 500 mM NaCl). The eluted fractions of the pure protein were desalted using an Econo-Pac 10DG (Bio-Rad, CA, USA) column pre-equilibrated with a desalting buffer containing 20 mM HEPES pH 7.5 and 300 mM NaCl. All the desalted fractions were pooled together and concentrated up to 5 - 8 mg/mL as determined by the Bradford assay using Bovine Serum Albumin (BSA) as a standard(Kruger, 2009). Concentrated protein fractions in small aliquots were stored at −80°C until further use.

### Purification of sub-methylated ribosome

*Thermus thermophilus* HB8 ΔKsgA strain (a gift from Dr. Hasan DeMirci, Stanford University, Palo Alto, CA, USA) was grown at 72°C in *Thermus* enhanced medium (American Type Culture Collection medium 1598) containing 25 μg/ml of Kanamycin. The cells were harvested and re-suspended in ribosome buffer (20 mM HEPES pH 7.5, 100 mM NH_4_Cl, 10 mM Mg(OAc)_2_, 6 mM 2-mercaptoethanol) and disrupted using sonication. After incubating the lysate with DNase I (10μg/ml) for 15 minutes, unlysed cells and cell debris were removed by centrifugation at 18000 rpm for 30 minutes at 4°C. Ribosomal particles were pelleted down by layering the supernatant on top of 30% w/v sucrose cushions made in a buffer containing 20 mM HEPES pH 7.5, 500 mM NH_4_Cl,10 mM Mg(OAc)_2_, 6 mM 2-mercaptoethanol and centrifuged with a 70Ti (Beckman coulter) rotor at 40,000 rpm for 18 hours at 4°C. The resulting glassy pellet was rinsed with ribosome buffer and re-suspended in 500 μl of the same buffer. 70S ribosomal particles were dialyzed in dissociation buffer (20 mM HEPES pH 7.5, 150 mM NH_4_Cl, 1 mM Mg(OAc)_2_, 6 mM 2-mercaptoethanol) to separate ribosome into its corresponding 50S and 30S subunits. The subunits were separated by zonal sedimentation on 13 ml 10 to 40% sucrose gradients (SW41 rotor at 40,000 rpm for 6 hours) in buffer (2 0 mM HEPES, pH 7.5, 50 mM NH_4_Cl, 1 mM Mg(OAc)_2_, 6 mM mercaptoethanol). The gradient was fractionated from top to bottom by pipetting 200 μl volume in separate vials. Each fraction was checked on agarose gel for its RNA integrity and subunit separation. The fractions containing pure 30S subunits were pooled together and dialyzed in a storage buffer (20 mM HEPES, pH 7.5, 150 mM NH_4_Cl, 1 mM Mg(OAc)_2_, 6 mM 2-mercaptoethanol) to remove sucrose from the sample. Purified fractions of 30S subunits were concentrated up to 4-5 μM as determined by standard conversion 1 A_260_ = 72pmol.

### Construction of 16S rRNA platform mutants

Mutation in 16S rRNA was introduced by PCR based overlap – extension protocol(Bryksin & Matsumura, 2010). The reaction mixture (50 μl) contained 1X GC phusion buffer, 0.2 mM deoxyribonucleotide triphosphate (all the PCR chemicals were supplied by Genetix Biotech Asia Pvt. Ltd., Mumbai, India), 1.6 μM of each of the primers, 1 ng/μl of template DNA and 1 U of Phusion polymerase (New England Biolabs). The mutant PCR fragments were then cloned into pSpurMS2 (a gift from Dr. Rachel Green, Johns Hopkins University) vector using KpnI and XbaI restriction sites, transformed into DH5α (for plasmid isolation) and the presence of the desired mutations was confirmed by nucleotide sequencing (Eurofins Genomics India Pvt., Ltd). Mutation driven structural change in RNA backbone was analyzed via RNA Composer, automated RNA structure 3D modeling server(Biesiada, Purzycka et al., 2016).

### Purification of MS2 fusion coat protein

The vector, pMAL– c2g containing MBP-MS2 (V75E A81G)-6X His (a gift from Dr. Gloria Culver, University of Rochester, Rochester, New York, USA) was transformed into *E. coli* expression strain BL21 (DE3) Gold. MS2 coat fusion protein purification was performed with a slightly modified version of the previously described procedure (Gupta & Culver, 2014). Cells were grown in Luria-Bertani (LB) medium containing 100 μg/ml ampicillin at 37°C until OD_600_ reached 0.6, overexpression of the protein was induced by 1 mM IPTG and grown further for 3 hours at the same temperature. Cells were harvested by centrifugation at 5000 rpm for 15 minutes and the pellet was re-suspended in lysis buffer (50 mM Tris-HCl (pH 7.5), 500 mM NaCl) and lysed by sonication. The lysate was incubated with DNase I (10 μg/ml, Thermo Scientific) and RNAse A (10μg/ml, Thermo Scientific) on ice for 10 minutes to reduce the co-purification of nucleic acid. Cellular debris in the lysate was removed by centrifugation at 18000 rpm for 30 minutes, before incubating with Ni-NTA resin pre - equilibrated with lysis buffer on a rocker for 1 hour at 4°C. The separated resin was loaded on a gravity column and washed with 200 ml of wash buffer (50 mM Tris-HCl (pH 7.5), 1 M NaCl, 50 mM Imidazole). The protein was eluted with elution buffer (50 mM Tris-HCl (pH 7.5), 500 mM NaCl, 500 mM Imidazole). The pure fraction of protein was pooled and dialyzed overnight in storage buffer (25 mM Tris-HCl (pH 7.5), 250 mM NaCl, 6 mM 2-mercaptoethanol, and 20% glycerol). Purified protein was stored at −80°C until further use.

### Affinity purification of MS2-tagged 30S ribosomal subunit

Untagged, unmethylated 30S subunits were purified from E. coli BW25113 ΔKsgA strain (a gift from Dr. Gloria Culver, University of Rochester, Rochester, New York, USA) via a standard technique of sucrose density gradient ultracentrifugation as described above. MS2 affinity purification system developed by Youngman and Green was used for obtaining pure, in vivo assembled 30S ribosomal subunits for our methylation assay (Youngman & Green, 2005). Ribosomes tagged with MS2 stem loop were expressed under the control of the lambda promoter. The plasmid pcI^857^ and either non-mutant or mutant pSpurMS2 transformed into Kasugamycin resistant *E. coli* BW25113 strain. pSpurMS2-WT and pSpurMS2-mutant were grown in 4×1L Luria-Bertani (LB) medium containing 100 μg/ml Ampicillin, 35 μg/ml Kanamycin, and 400 μg/ml Kasugamycin at 42°C until OD_600_ reached ~ 0.6. The culture was chilled for 1 hour to obtain runout ribosomes before cells were harvested at 5000 rpm for 15 minutes. Total crude ribosome was purified from the lysate using sucrose cushion as previously discussed. The ribosome pellet was washed with buffer (20 mM Tris-HCl (pH 7.5), 100 mM NH_4_Cl, 1 mM Mg(OAc)_2_, 5 mM 2-mercaptoethanol) and re-suspended in the same buffer. 10 mg of MBP-MS2-His protein were incubated with Ni-NTA resin pre-equilibrated with MS2 fusion coat protein binding buffer (25 mM Tris-HCl (pH 7.5), 250 mM NaCl) by shaking at 4°C for 15 minutes, and then washed with binding buffer to removed unbound protein. Separated fusion protein bound resin was mixed with 10 ml of the crude ribosome (100 mg/ml) in a 50 ml disposable falcon tube on a rotatory platform at 4°C for 30 minutes. Complex bound to resin was loaded on a gravity column and washed with buffer containing 20 mM Tris-HCl (pH 7.5), 200 mM NH_4_Cl, 1 mM Mg(OAc)_2_, 5 mM 2-mercaptoethanol, and the tagged ribosomes were eluted with elution buffer (20 mM Tris-HCl (pH 7.5), 100 mM NH_4_Cl, 1 mM Mg(OAc)_2_, 5 mM 2-mercaptoethanol, 300 mM Imidazole). Eluted tagged mutant 30S subunits were dialyzed (20 mM Tris-HCl (pH 7.5), 100 mM NH_4_Cl, 1 mM Mg(OAc)_2_, 5 mM 2-mercaptoethanol) and concentrated up to 1 nmol/ml (1A_260_ = 69 pmol) using Amicon Ultra centrifugal filters (Millipore, MWCO 100,000). Further, its integrity was checked on 10–40% sucrose gradient and on RNA gel. Pure tagged mutant 30S subunit was stored at −80°C in aliquots for in vitro methylation assay.

### Scintillation Assay

The in vitro methylation assay was adapted from O’Farrell et al (O’Farrell, Pulicherla et al., 2006). The assay was performed in a methylation buffer (50 mM Tris-HC l (pH 7.5), 40 mM NH_4_Cl, 4 mM Mg(OAc)_2_, 6 mM 2-mercaptoethanol) containing 0.5 μM of RNA, 0.5 μM of methyltransferase, 0.1 μM (3H)-S-adenosyl-L-methionine (SAM) [(methyl-3H) AdoMet, 100 Ci/mmol, ARC], and 1U of RNase inhibitor (Thermo Scientific) in total reaction volume of 50 μl. Reactants were mixed and incubated at 37°C for 30 minutes in the buffer described above. After 30 minutes, pre-chilled 10 μl of 100 mM unlabeled SAM (Sigma-Aldrich) was added, to quench the reaction. The quenched reaction was deposited on pre-charged filter paper (Nylon 66 filter membrane), washed thrice with chilled 5% TCA to remove the non-specific binding, and rinsed briefly with 100% ice-cold ethanol. The filter paper was air dried for 1 hour and then placed into 2 ml of scintillation liquid (Ultima Gold, PerkinElmer), and radioactivity was counted using a scintillation counter (Tri-Carb B2810TR; PerkinElmer, USA).

### Growth analysis of pSpurMS2 plasmids containing platform mutation

Construction of Δ7 prrn strain exclusively expressing mutant ribosome requires shuffling of Ampicillin (Amp) resistant wild type and mutant pSpurMS2 plasmid with the wild type rrn pCsacB7 plasmid(Sun, Vila-Sanjurjo et al., 2011). MC338 strain (a gift from Dr. Gloria Culver, University of Rochester, Rochester, New York, USA) lacks chromosomal rDNA and supports the growth of cells by plasmid pCsacB7, which carries wild type rrnC operon, Kanamycin resistance marker, and SacB gene, conferring sensitivity to sucrose (Sun et al., 2011). The wild type and mutant pSpurMS2 Amp resistant plasmid were transformed in MC338 strain, single Amp resistance colony was grown in LB media and subjected to sucrose selection on a sucrose-Amp plate. The sucrose resistant plasmid was then tested for loss of Kanamycin resistance. The inability of certain mutant (Amp resistant) plasmid to replace wild type Kanamycin resistant plasmid (observed by the persistence of Kanamycin resistance after selection on a sucrose-Amp plate) was confirmation of rRNA deleterious mutation, to the extent Amp resistant cells harboring mutant pSpurMS2 plasmid were grown in LB media at 37°C overnight. The saturated culture was sub cultured to an OD_600_ of 0.1 in LB media containing 100 μg/ml of Ampicillin. Cells were incubated with shaking (225 rpm) at 37°C and OD_600_ was monitored. The experiment was done in triplicate and data was plotted on a logarithmic scale and fitted with equation Y = Ae^BX^. The doubling time (T_d_) was calculated by using equation T_d_ = ln(2)/B.

### Co-sedimentation Assay

The complex was prepared as described previously by Boehringer *et al*. (Boehringer et al., 2012). The unmethylated 30S ribosomal subunit was purified as described above. The 30S-KsgA complex was prepared by incubating KsgA (20- and 50-fold excess) with 30S ribosomal subunits (10 pmol) for 15 minutes at 37°C in buffer A (50 mM HEPES (pH 7.5), 40 mM NH_4_Cl, 4 mM Mg(OAc)_2_ and 6 mM 2-mercaptoethanol). Reactions were quenched by incubation on ice for 10 minutes and quickly spun at 10000 rpm. The reaction was loaded on top of 150 μl of 30% sucrose cushion in buffer A and centrifuged at 100,000 x g for 60 minutes. Complex formation was checked by running supernatant and pellet fractions, on 15% SDS – PAGE and stained with Coomassie blue (Fig S1).

### Statistical analysis

The data from all the three experimental replicates were presented as mean ± standard deviations. Differences between the groups were evaluated using Student’s t-tests. P values <0.05 were considered significant. Statistical significance in Figs is displayed as for ** P < 0.01, for *** P < 0.001.

### Preparation of the KsgA-30S Complex

The complex was reconstituted by incubating purified sub-methylated 30S ribosomal subunit (2μM final concentration) with KsgA at a stoichiometric ratio of 1:20. The reaction was set up in reconstitution buffer (50 mM HEPES (pH 7.5), 40 mM NH_4_Cl, 4 mM Mg(OAc)_2_, 6 mM 2-mercaptoethanol) at 37 °C for 15 minutes followed by a quick spin at 2000 x g and incubation on ice for 90 minutes. The 50 μl reaction was diluted to 1 ml by adding reconstitution buffer and the complex was pelleted by ultracentrifugation at 55000 rpm in TLA55 rotor for 15 hours at 4°C. The supernatant was discarded and the pellet was re-suspended in 15-20 μl of reconstitution buffer followed by a 2 minutes spin at 500 x g to remove any debris. The supernatant was aliquoted carefully into a new vial and A_260_ was measured with an spectrophotometer.

### Grid preparation and Cryo-EM data collection

The reconstituted 30S-KsgA complex (3 μL) was applied to a glow discharged Quantifoil holey carbon grid (R 1.2/1.3, Au 300 mesh) at a concentration of 2 μM on a Vitrobot (FEI) set at 4 °C and 100% humidity with 15 second incubation and 3 - 3.5 second blot time. The grids were screened for good quality ice, particle distribution, and density before setting up data collection. Data were collected on a Titan Krios G3 using the Falcon 3 detector in integration mode at a sampling rate of 1.38 Å (nominal magnification of 59000X and calibrated magnification of 101449) and an exposure of 2 seconds with a total dose of 43.06 e/Å2 fractionated over 30 frames (1.43 e/Å2 per frame). A defocus range of −1.8 to −3.5 was used and 2154 movies were collected.

### Data processing, class reconstruction, and model building

Image processing was done using Relion (3.0 and 3.1) (Zivanov, Nakane et al., 2018, Zivanov, Nakane et al., 2020). Movies were motion corrected by implementing Relion’s inbuilt algorithm using a 5 x 5 patch configuration (Scheres, 2014, Zheng, Palovcak et al., 2017). The contrast transfer function (CTF) of the aligned micrographs was estimated with Gctf 1.06(Zhang, 2016). Particles were picked using Gautomatch (http://www.mrc-lmb.cam.ac.uk/kzhang/) using templates from previously collected data. A total of 706511 particles were extracted using a box size of 320 pixels and 2D classified into 200 classes using a mask diameter of 280 Å. Bad classes were purged and classes with good particles were used for 3D refinement followed by 3D classification (10 classes) with resolution limited to 8 Å. The 3D classes were checked in Chimera and the classes with broken/damaged particles were discarded.

A total of five classes were 3D refined and post-processed (Fig S2). The particles from 3D refined classes were used for estimation of per-particle defocus, beam-tilt, aberration, and anisotropic magnification estimation followed by refinement and Bayesian polishing (Zivanov, Nakane et al., 2019). The B-factor weighted particles were further subjected for 3D refinement. All classes were subjected to another round of CtfRefinement and 3D refinement. The resulting reconstructions were used for post-processing and local resolution estimation (LocRes). The reconstructions of all classes were used for multi-body refinement using 2 body (body 1 = 30S Body + Platform + KsgA and body 2 = 30S head) or 3 body architecture (body 1 = 30S Body + Platform, body 2 = 30S head and body 3=KsgA+h44+h27+h24). In classes that had disordered h44 in reconstructions, body 3 did not contain h44. The 3 body multi-body refinement was successful in capturing the KsgA movement leading to improved maps. There was an improvement in the 30S head and body also leading to a better capture of the h44 movement. The multibody refinement maps were combined using Combine Focused Maps (Phenix) (Adams, Afonine et al., 2010). These maps with improved resolution for head, body, and KsgA region were used for model fitting and building (Fig S3, S4). In addition, we also used the blurring option in Coot(Emsley, Lohkamp et al., 2010) (Cryo-EM module) for the post process (both for the combined or multibody) maps, local_aniso_sharpen within Phenix (with the combined unsharpened map and the model without description of resolution), and the maps using LocSpiral (Kaur, Gomez-Blanco et al., 2021) to aid in model building.

The class K1 was sub-classified into four classes yielding two classes in apo (K1-k2) and ksgA bound (K1-k4) forms. These two classes were also subjected to multi-body refinement using the 3 body architecture. Model fitting and building were done in Chimera and COOT (Emsley et al., 2010, Pettersen, Goddard et al., 2004).We used *Thermus thermophilus* (Tt) HB8 30S ribosome pdb (PDB-4B3R) for preliminary fitting onto the 30S (class K1-k2) followed by manual fitting in COOT(Emsley et al., 2010). The model was built by iterations in COOT and Phenix real space refinement and Refmac (Adams et al., 2010, Afonine, Poon et al., 2018, Emsley et al., 2010, Murshudov, Skubak et al., 2011). Density for KsgA was fitted with Bacillus subtilis KsgA structure (PDB-6IFS) in Chimera followed by model building and fitting in Coot. Various regions of the 30S-KsgA complex where density was absent like h44 region, CDR, N-terminal loop of KsgA etc., were deleted from the model. There are additional potential densities for Mg^2+^ ions but to be consistent across all classes and models, these have not been modeled. The models for other classes (K1-k4, K2, K4, K5, and K6) were generated using the coordinates from K1-k2 taking into account the head movements and absence of density in the map. The refined models were validated with MolProbity within Phenix (Chen, Arendall et al., 2010). Figures were prepared using UCSF Chimera, ChimeraX (Goddard, Huang et al., 2018, Pettersen et al., 2004) and Pymol 2.4 (The PyMOL Molecular Graphics System, Version 1.2r3pre, Schrödinger, LLC).

## DATA AVAILABILITY

CryoEM maps and coordinates have been deposited in PDB and EMDB under following codes: Class K1k2 – 7V2L & EMD-31655; Class K1k4 - 7V2M & EMD-31656; Class K2 - 7V2N & EMD-31657; Class K4 - 7V2O & EMD-31658; Class K5 - 7V2P & EMD-31659; Class K6 - 7V2Q & EMD-31660

## ACKNOWLEDGEMENT

The authors acknowledge the National CryoEM Facility at Bangalore Life Science Cluster for EM data collection and processing and Scintillation facility at IIT Bombay. We thanks Prof. Gloria Culver (University of Rochester) and Prof. Rachel Green (Johns Hopkins University) for sharing strains and plasmid. JS thanks University Grants Commission (UGC) and IIT Bombay for PhD fellowship. RA acknowledges DBT/Wellcome Trust-India Alliance Fellowship [grant number IA/S/19/1/504293] and STARS [grant number MoE/STARS-1/APR2019/BS/523/FS]. KRV acknowledges SERB, India for the Ramanujan Fellowship (RJN-094/2017), DBT B-Life grant DBT/PR12422/MED/31/287/2014 and the support of Department of Atomic Energy, Government of India, under Project Identification No. RTI4006.

## Author Contributions

RA and JS designed and performed all the experiments. JS, RR and KRV performed CryoEM work. JS performed all the biochemical experiments under supervision of RA. All authors have contributed equally in the manuscript. JS and RR are considered first author.

## Supporting Information

**Figure S1.**
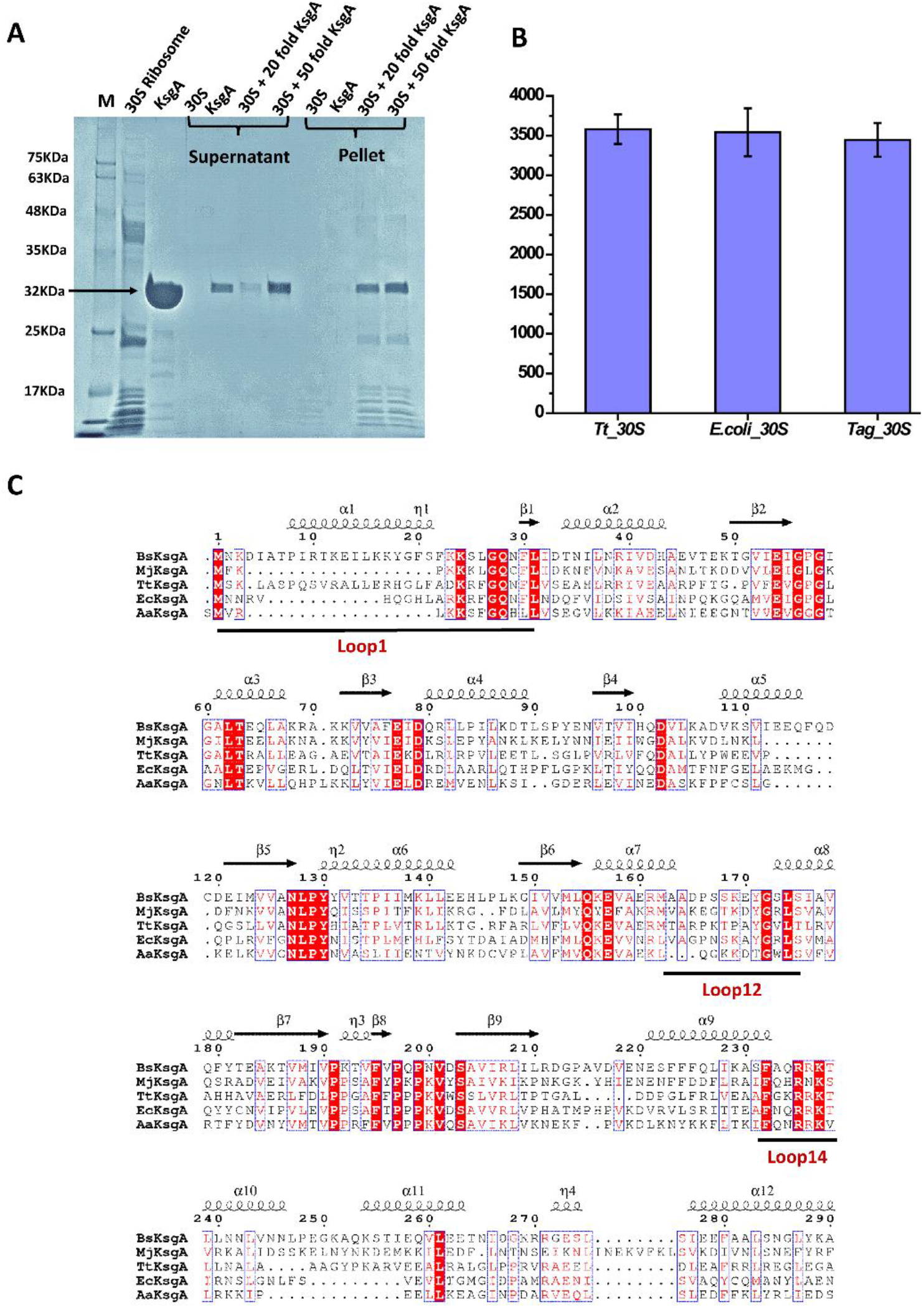
Binding of KsgA with 30S ribosomal subunit. **(A)** SDS-PAGE showing the binding of B. subtilis (Bs) KsgA with T. thermophilus (Tt) 30S subunit. Control KsgA and 30S has been loaded in high concentration (30μM KsgA, 25 pmol 30S). **(B)** In-vitro scintillation assay of T. thermophilus, E.coli, MS2 tag-E.coli 30S subunit with BsKsgA. **(C)** Secondary structures of B. subtilis (Bs) KsgA and sequence alignment with other species. The plot was generated using the ESPript 3.0 server (Robert et al., 2014).

**Figure S2.**
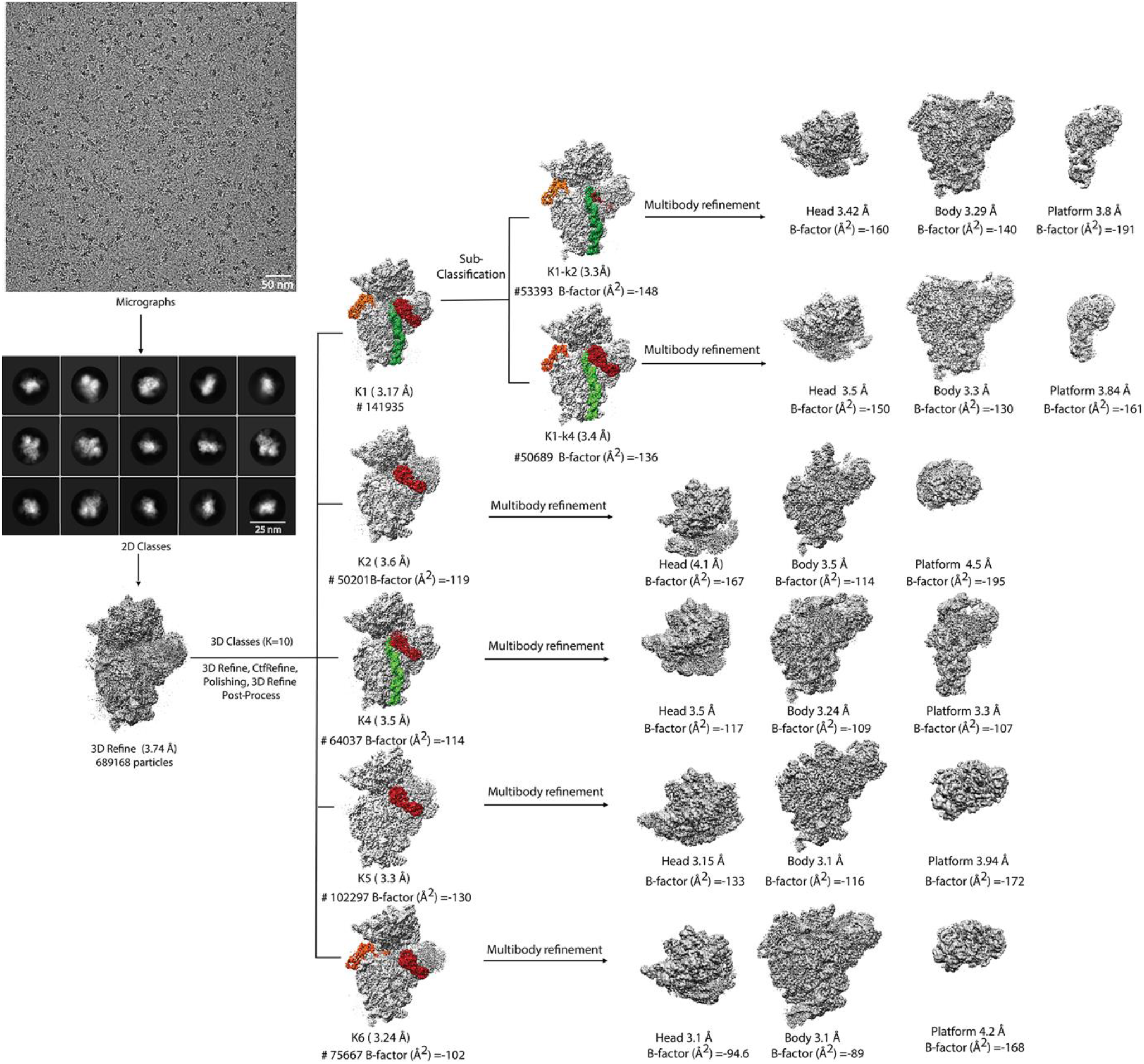
Cryo-EM data processing work flow of the 30S-KsgA complex. The flow chart shows the processing steps and relevant statistics for the data collection of 30S-KsgA. Based on inspection of the maps from 3D classification, five classes were selected for further processing. Class K1 was further classified into four classes and two classes (K1-k2 and K1-k4) were selected from these four classes. The B-factor sharpened maps for all these classes are shown in the flowchart with the h44 segment colored in green, KsgA interacting at P1 and P2 in red and orange. All these classes were subjected to multibody refinement using 3 body setup for head, body and platform/KsgA regions. These maps were used in model building, particularly at the KsgA/platform interface.

**Figure S3.**
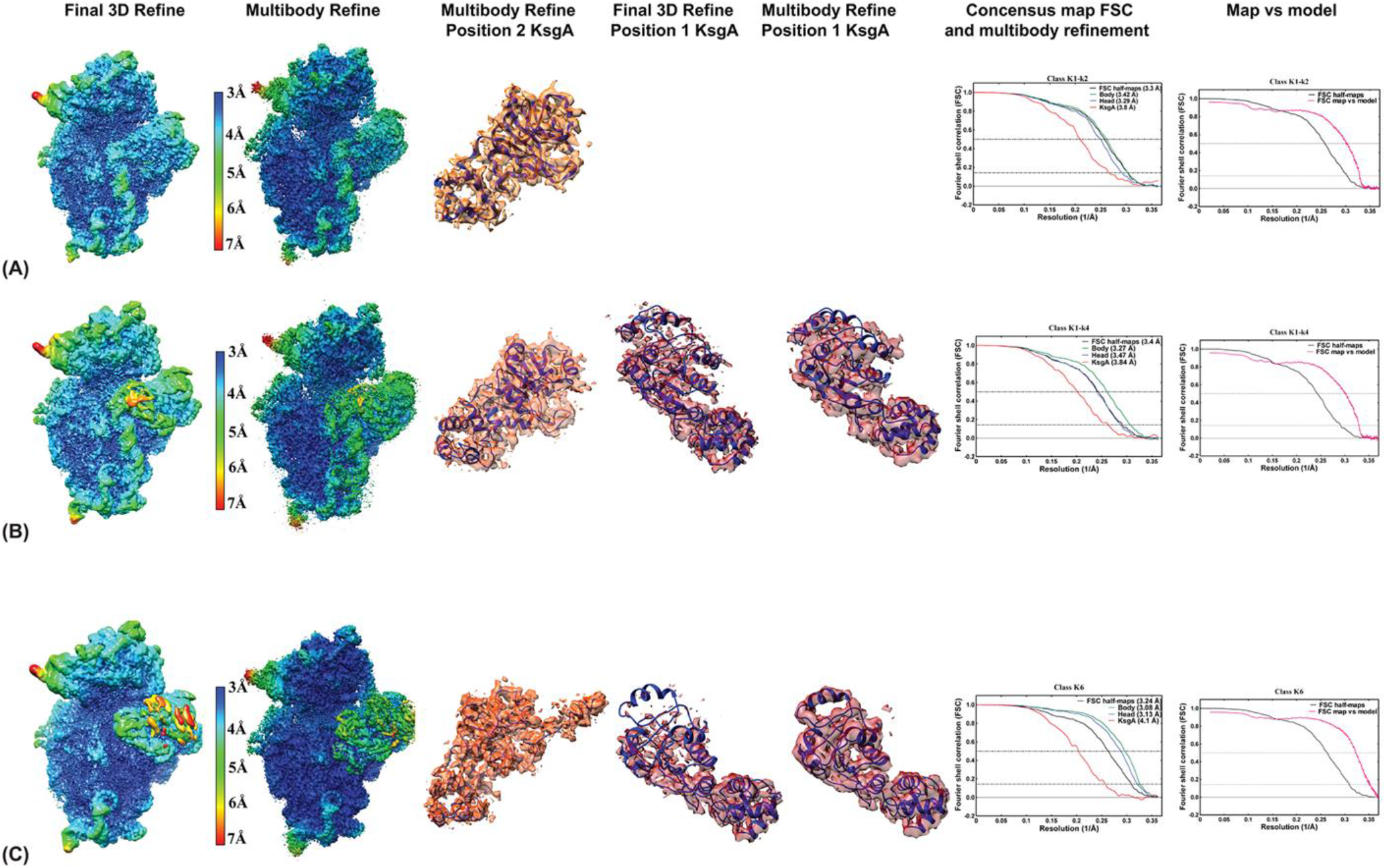
Refinement of classes K1–k2 (A), K1-k4 (B) and K6 (C). The figure shows local resolution maps for the postprocess maps from the 3D refinement of the classes in the left most column. The second column shows the local resolution maps for the postprocess maps of the multibody refinement. Improvement in the resolution are seen from the color key. The class K1-k2 has very little ordered density for KsgA at the platform region pointing towards the initial step of KsgA binding with 30S. The figure in column 3 shows the maps for the position 2 KsgA near helix 16. Column 4 and 5 shows improvement of map for platform KsgA after multibody refinement. The figures for the map were made in Chimera using the composite map. The sixth and seventh column show FSC curves of head, body and platform regions with respect to 3D refinement maps and the map vs model respectively.

**Figure S4.**
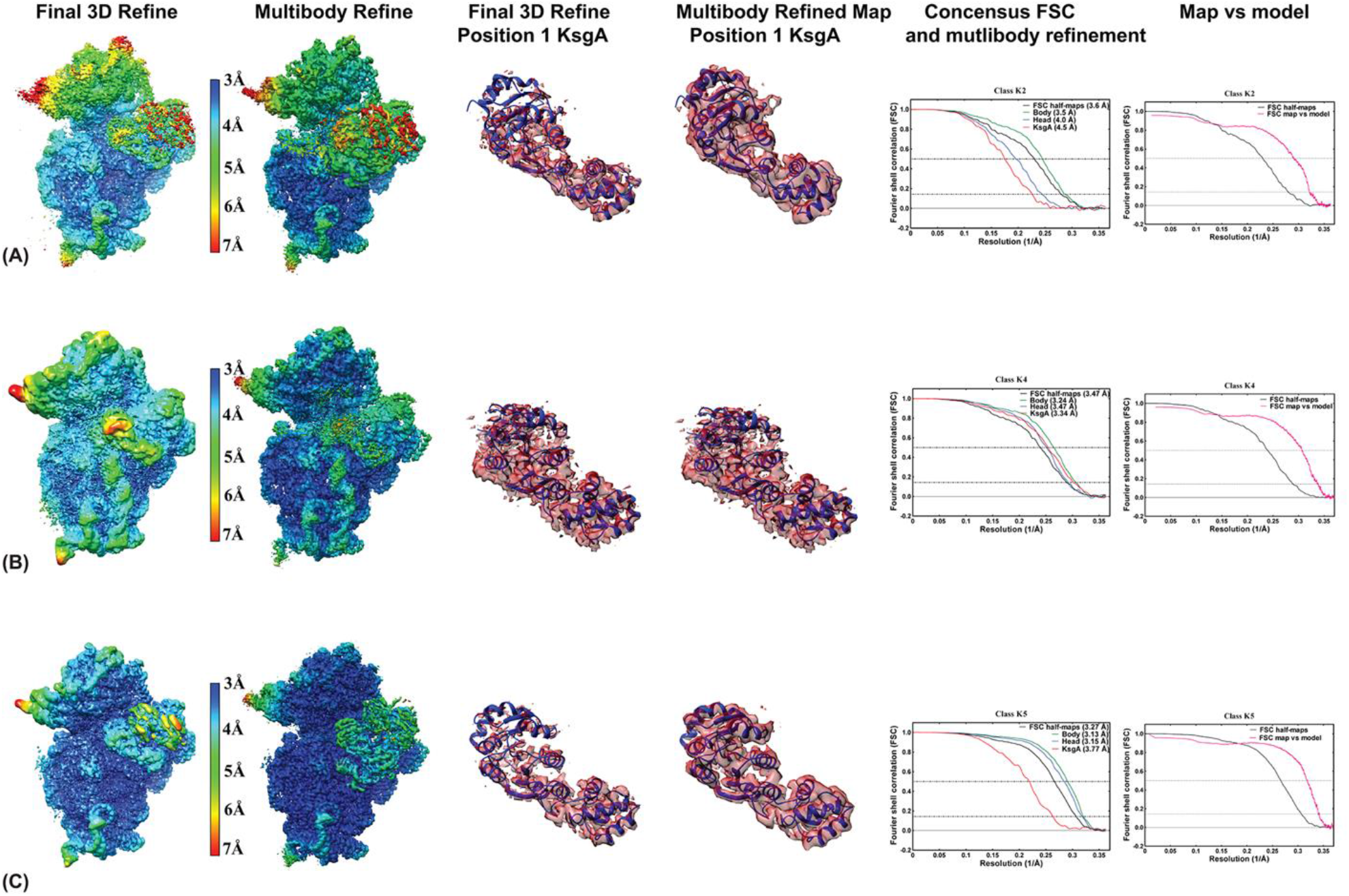
Refinement of class K2 (A), K4 (B) and K5 (C). Local resolution maps for the postprocess maps from the 3D refinement of the classes are shown in the left most column along with the local resolution map for the postprocess maps of the multibody refinement in the second column. The map shows improvement in resolution as seen from the color key. Comparison of column 3 and 4 of A, B and C show the improvement of the map for platform KsgA after multibody refinement. The figures for the map were made in Chimera using the composite map. The fifth and sixth column shows the FSC curves of head, body and platform regions with respect to 3D refinement maps and the map vs model respectively.

**Figure S5.**
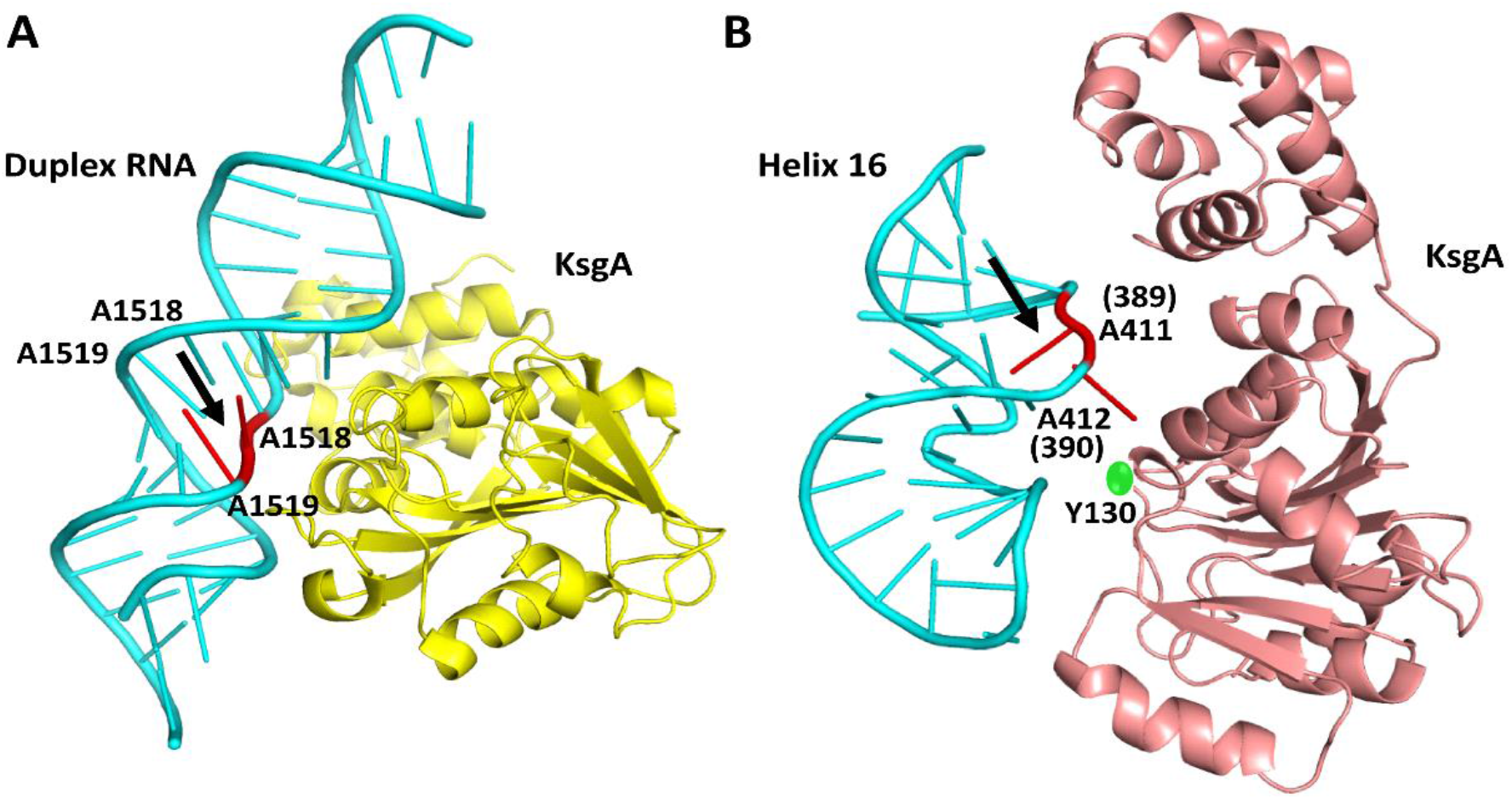
Comparison of RNA binding mode of KsgA. **(A)** A. aeolicus KsgA (PDB ID 3FTF, yellow) shows N-terminal domain is interacting with RNA near to the target residue (red) (Chao Tu et al., 2009). **(B)** Position 2, BsKsgA interacting with h16, sequence and structure resembles to h45 tetraloop which resulted in spurious binding. Active site Y130 is highlighted in green sphere to show the distance between A412 (Thermus thermophilus numbering shown in bracket) and active site of the enzyme highlighting that the RNA is not positioned close enough to KsgA for catalysis.

**Figure S6.**
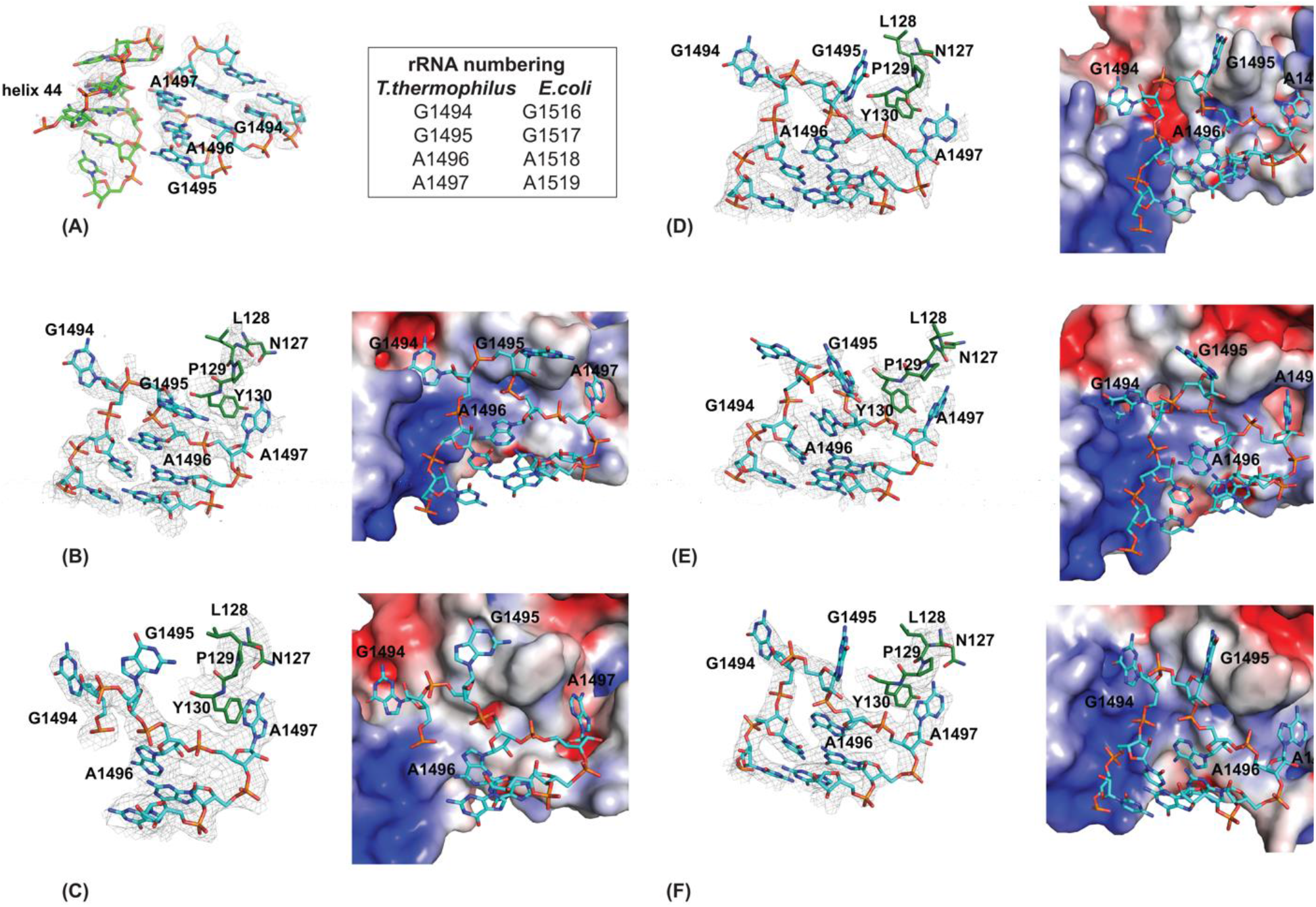
Adenosine dimethylation site for all the six classes showing the EM map and the electrostatic surface view of KsgA with RNA. **(A)** shows the EM map for h44 and h45 for the K1-k2 class where KsgA density is very weak and not modelled. **(B, C, D, E &F)** shows the EM map for the KsgA active site (NLPY) in proximity with 30S dimethylation site (AA:1496-1497; E. coli and T.thermophilus numbering are shown in the box) in classes K1-k4, K6, K2, K4, and K5 and the electrostatic surface view for KsgA with h45 loop in the positively charged grooves.

**Figure S7.**
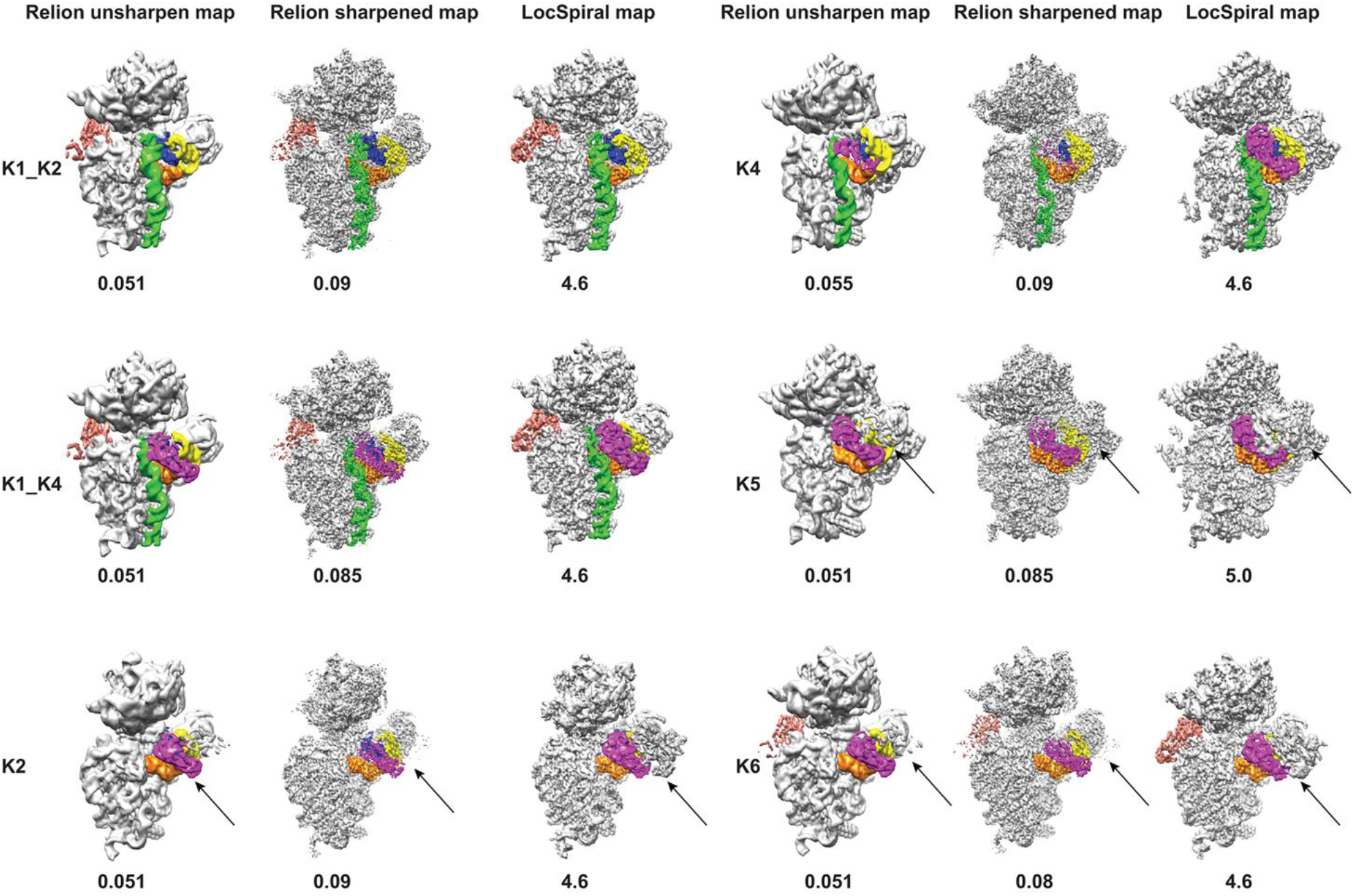
CryoEM maps of the classes that were used in model building. For each class, the unsharpened, combined map from Relion 3D refine, sharpened map from Relion postprocess procedure and the LocSpiral map are shown. The numbers below each map are the threshold (in Chimera) used to depict the map. In the case of LocSpiral maps, hide dust option was used. The RNA helices h44 (green), h45 (blue), h24 (yellow) and h27 (orange) are shown in each map. KsgA molecules in P1 and P2 positions are colored in magenta and salmon respectively. The arrow mark in classes K2, K5 and K6 mark the density for additional KsgA molecules that are present but not modelled. These maps are shown to highlight the blurring (or fragmenting) of regions of the maps that are at lower resolutions (example KsgA) and the use of multiple maps for model building. To show the extra density near P1 of KsgA, the maps have been slightly rotated for classes K2/K6.

**Figure S8:**
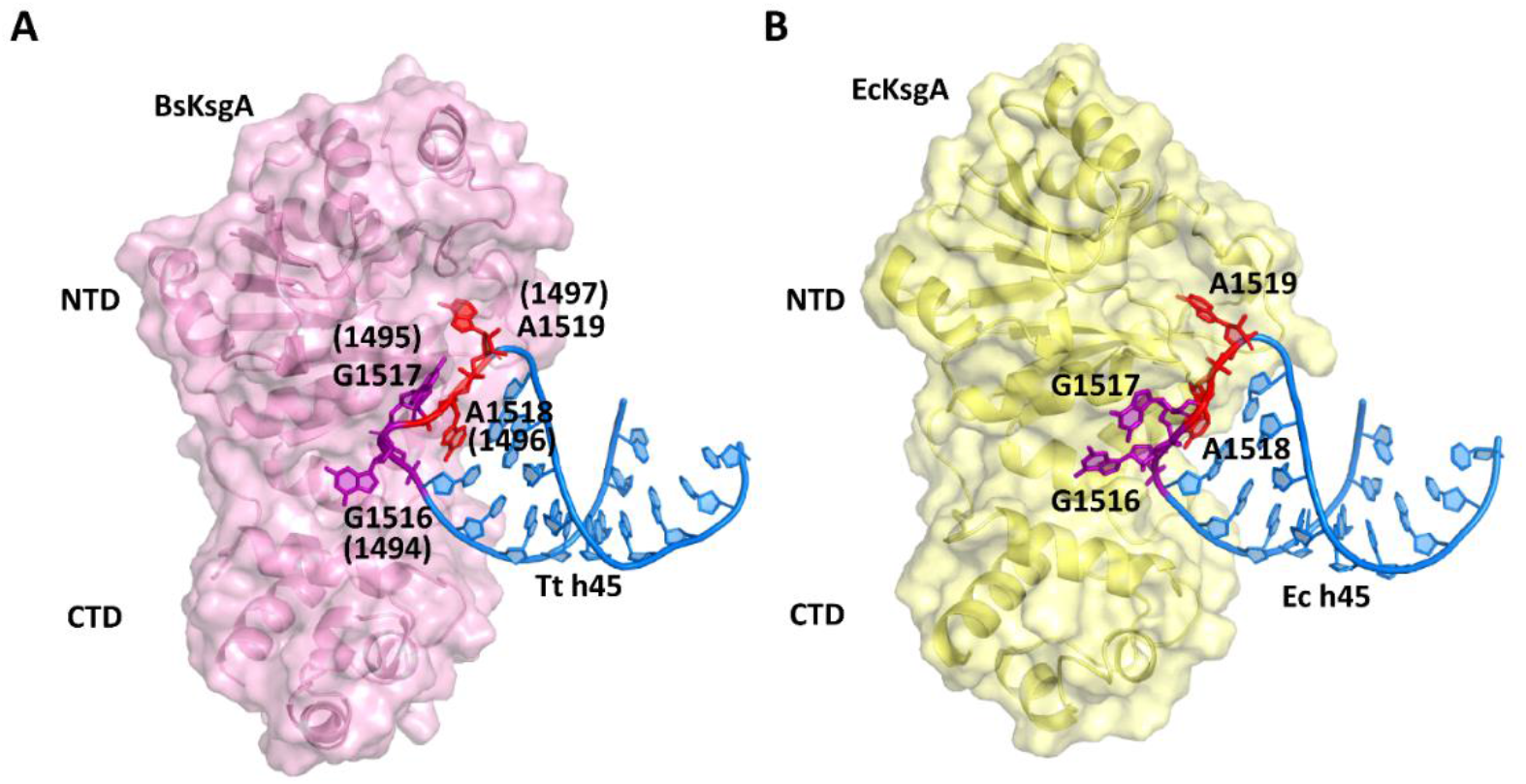
Comparison of binding modes of KsgA and h45. **(A)** BsKsgA (pink) (this study) **(B)** EcKsgA (pale yellow) (PDB ID: 7O5H) (Stephan et al., 2021). BsKsgA interacting with Tt h45 (marine blue) with target base A1519 (red) flipped inside the catalytic pocket, whereas the terminal base G1516 (purple) is flipped inside the interface exclusive pocket. Other tetraloop base G1517 (purple) is dynamic in nature, second target base A1518 (red) resides inside the tertraloop preparing for the subsequent methylation event. The same interaction for EcKSgA with Ec h45 is shown in panel B.

**Figure S9:**
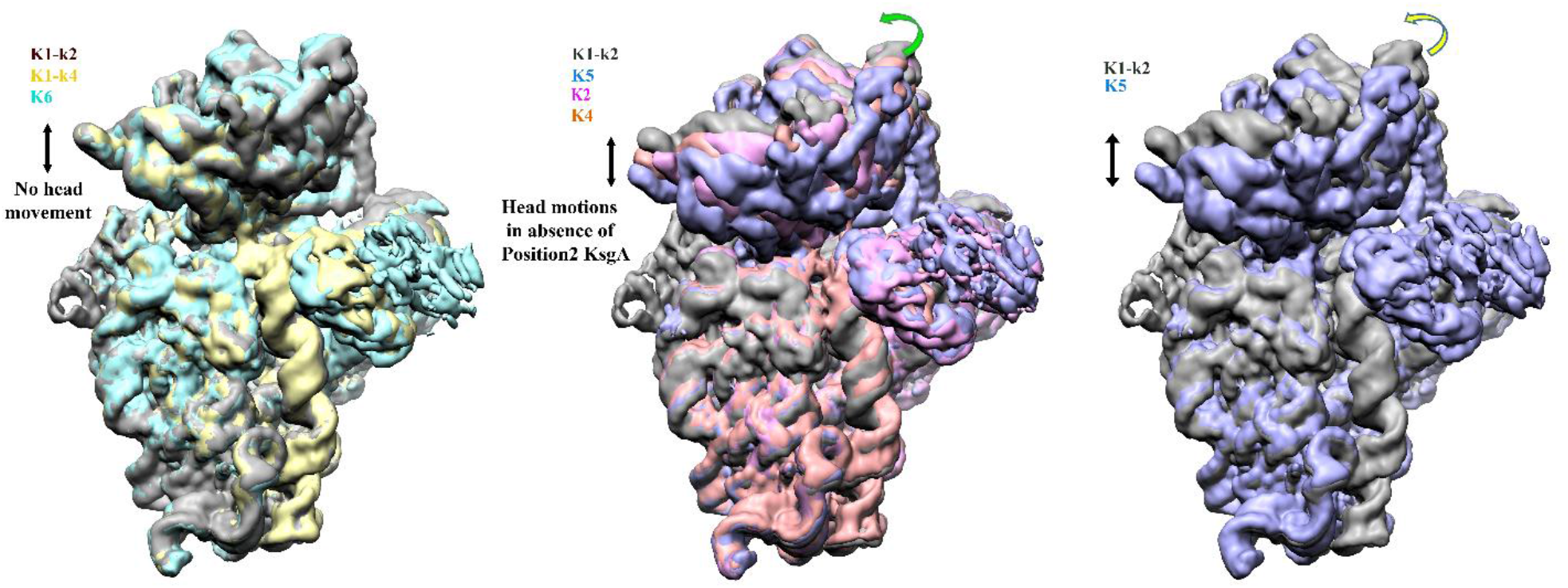
Comparison of head positions in all six classes. The left figure shows the comparison of K1-k2, K1-k4, K6 where an extra density is present near helix 16. This creates a lock for the head domain fixing it in a non-latched mRNA positions. The middle figure shows the map of K2, K5, K4 superposed with K1-k2. The head shows forward motion towards the body of the 30S in classes with KsgA absent near helix 16. The degree of freedom is very clear in the figure on right where K5 is compared with K1-k2.

**Figure S10.**
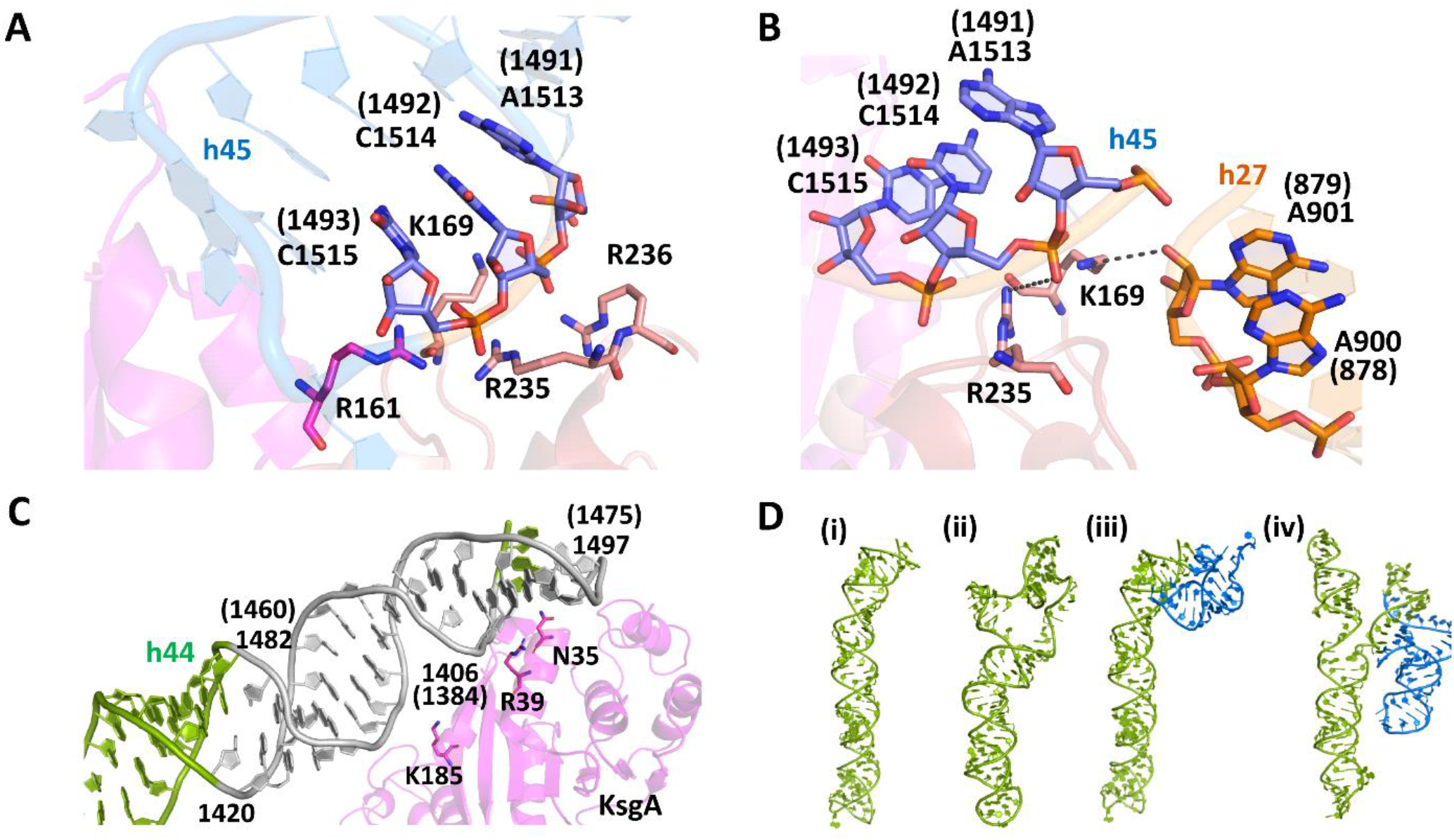
Significance of helix 44 in 30S ribosomal subunit. **(A)** Electrostatic interaction between stem residues (C1515, C1514) of helix 45 with positively charged residues (R161, K169, R235, R236) of KsgA. **(B)** K169 in KsgA loop12, K169 can form potential hydrogen bonds with h45 and A901 (h27). R235 from loop14 interacts with h45 phosphate backbone. **(C)** Interaction of h44 with KsgA, rRNA region shown in gray showed protection in chemical modification experiment (Zhili Xu et al., 2008). **(D)** Conformational rearrangement in h44 induced by mutagenesis as predicted by RNA folding analysis by RNA Composer wild type h44 (ii) h44_LM construct (iii) wild type h44 and h45 arrangement in 30S (iv) h44 and h45 arrangement in h44-LM mutant.

**Figure S11.**
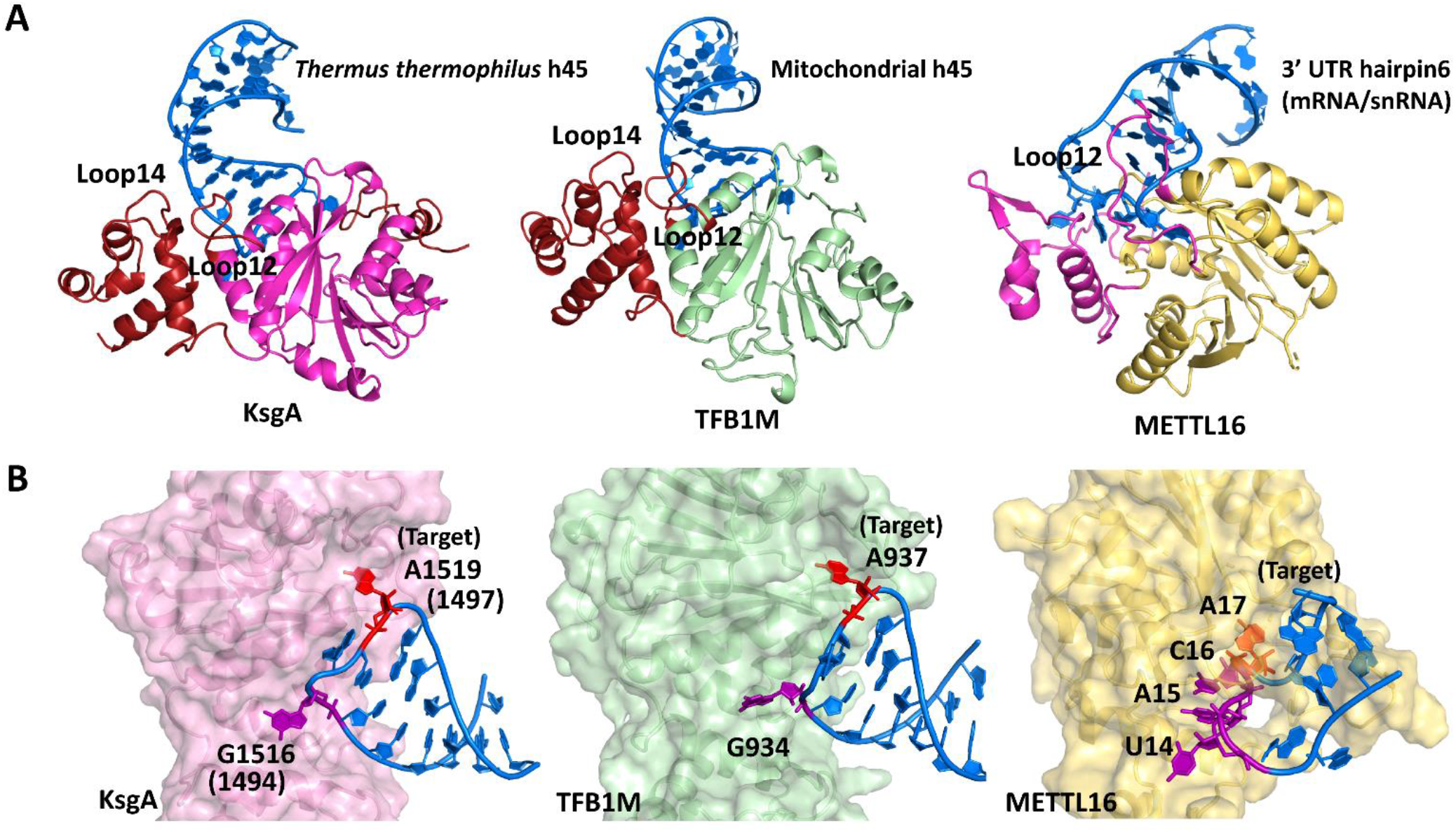
Comparison of conformation of the target helix for different rMTases. **(A)** In the left panel shows the current structure of KsgA (magenta, loop12 and C-domain highlighted with firebrick red) in complex with h45 tetraloop of 30S. The middle panel shows the structure of human mitochondrial ribosomal MTase, TFB1M (palegreen, loop12 and C-domain highlighted with firebrick red) in complex with RNA stemloop (PDB ID: 6AAX). The right panel shows the structure of mRNA/snRNA MTase METTL16 (yelloworange, loop12 and C-domain highlighted with magenta) in complex with RNA hairpin (PDB ID: 6DU5) **(B)** Zoom view of target recognition by MTase. A dual base flipping is evident in both KsgA and TFB1M, whereas multiple base flips are visualized in METTL16 hinting that base flipping at subsidiary sites is crucial for recognition.

**Figure S12.**
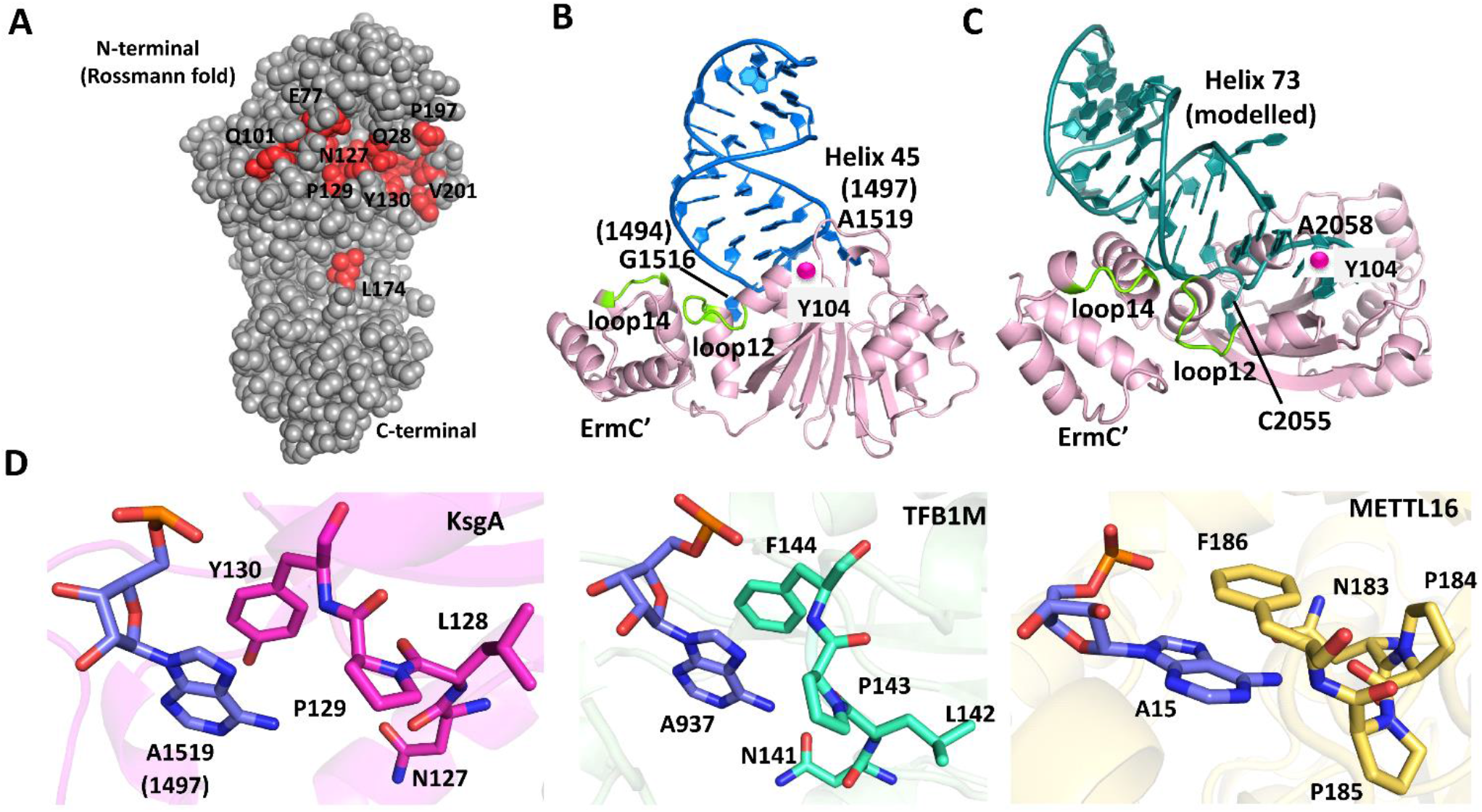
Comparison of RNA MTase. **(A)** Conserved residues of Rossmann fold are shown in red and variable residues are shown in gray. **(B)** To analyse the specificity targeting of loop12 and loop14, ErmC’ is superimposed on KsgA. It is shown that ErmC’ loops, shown in green are not able to stabilize the KsgA substrate i.e., h45. **(C)** Coordinates of h73, where ErmC’ methylates were extracted from PDB ID: 2AVY and superimposed on h45-KsgA complex, which shows the binding mode of A2058 with ErmC’. Similar to KsgA, the residue C2055, 15 Å away likely flips into a distal pocket. **(D)** Active site comparison of RNA methyltransferases.

**Table S1.**
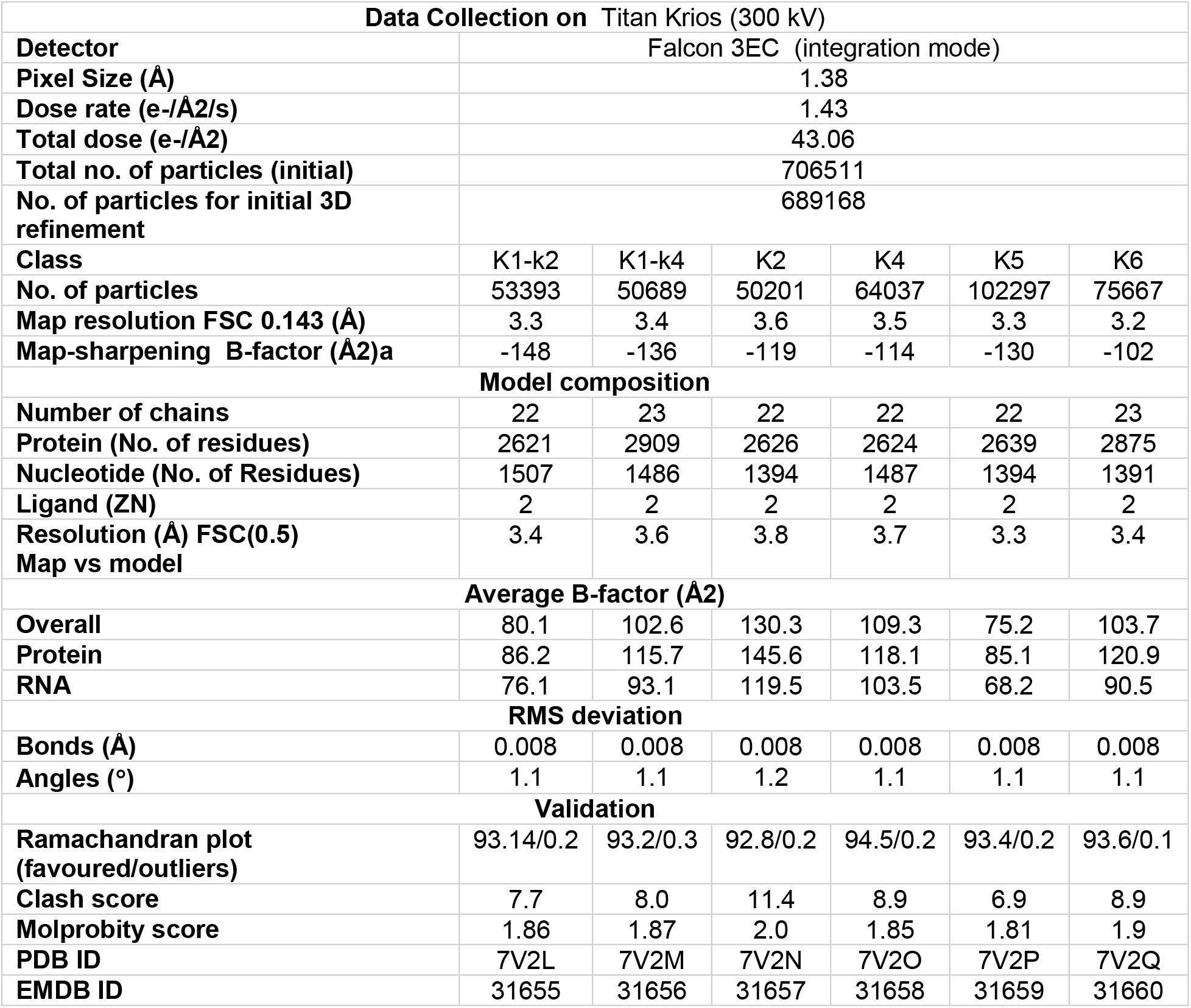
Cryo-EM Data collection and refinement statistics for KsgA-30S complexes. a - The map-sharpening B-factor (Å2) are the values from the postprocess job for each class after 3D refine. Multiple maps as described in methods were used for model building.

**Table S2.**
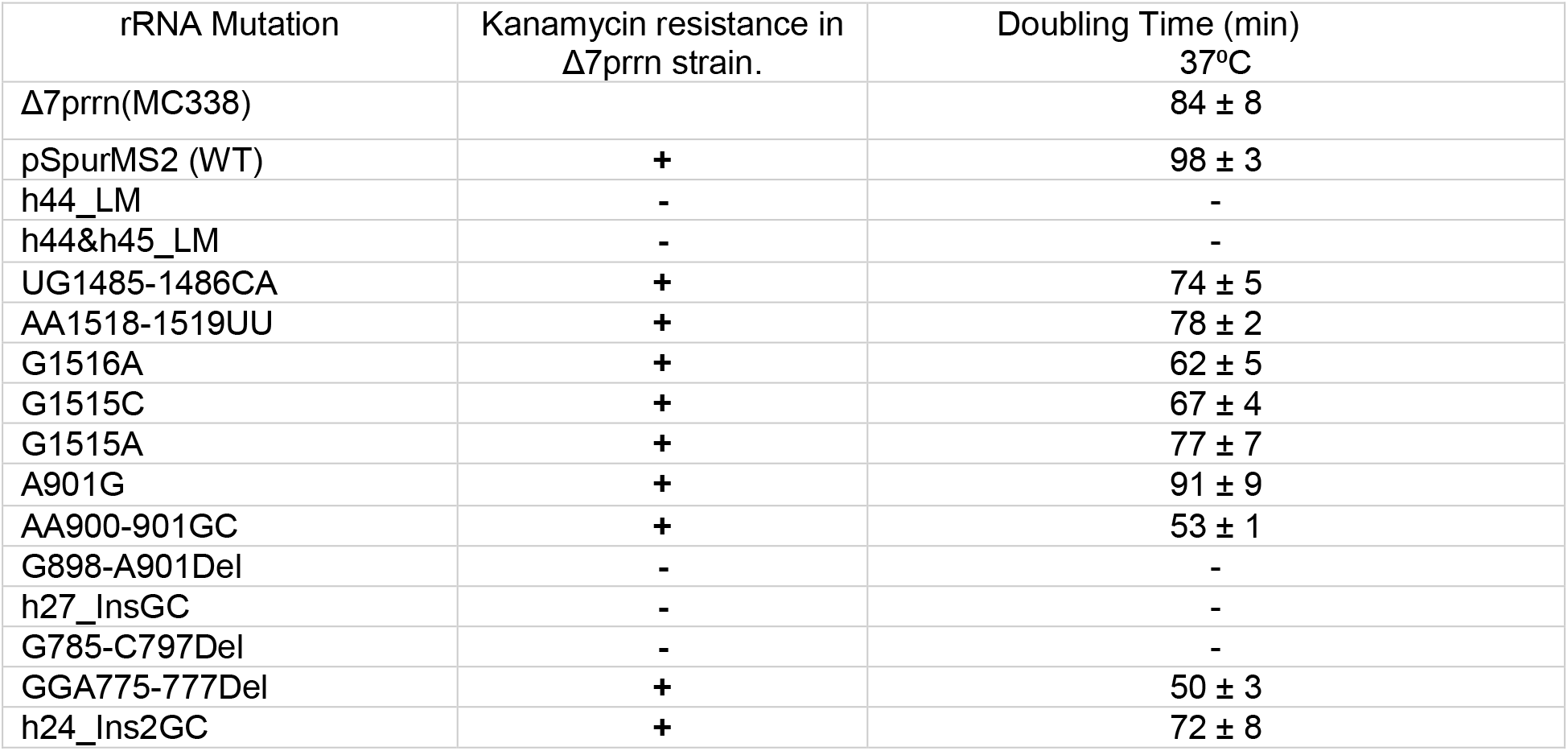
Growth characteristics of 16S rRNA mutants in MC338 strain

